# Covalent Aurora A regulation by the metabolic integrator coenzyme A

**DOI:** 10.1101/469585

**Authors:** Yugo Tsuchiya, Dominic P Byrne, Selena G Burgess, Jenny Bormann, Jovana Bakovic, Yueyan Huang, Alexander Zhyvoloup, Sew Peak-Chew, Trang Tran, Fiona Bellany, Alethea Tabor, AW Edith Chan, Lalitha Guruprasad, Oleg Garifulin, Valeriy Filonenko, Samantha Ferries, Claire E Eyers, John Carroll, Mark Skehel, Richard Bayliss, Patrick A Eyers, Ivan Gout

## Abstract

Aurora A is a cell cycle protein kinase implicated in multiple human cancers, and several Aurora A-specific kinase inhibitors have progressed into clinical trials. In this study, we report structural and cellular analysis of a novel biochemical mode of Aurora A inhibition, which occurs through reversible covalent interaction with the universal metabolic integrator coenzyme A (CoA). Mechanistically, the CoA 3’-phospho ADP moiety interacts with Thr 217, an Aurora A selectivity filter, which permits the formation of an unprecedented covalent bond with Cys 290 in the kinase activation segment, lying some 15 Å away. CoA modification (CoAlation) of endogenous Aurora A is rapidly induced by oxidative stresses at Cys 290 in human cells, and microinjection of CoA into mouse embryos perturbs meitoic spindle formation and chromosome alignment. Aurora A regulation by CoA reveals how targeting of Aurora A might be accomplished in the future by development of a ‘double-anchored’ covalent inhibitor.

## Introduction

Aurora kinases are Ser/Thr kinases that play well-documented roles in eukaryotes, where they control meiosis, mitosis and cell division (Carmena and Earnshaw, 2003). Aurora kinases differ in their relative expression levels, stability and sub-cellular localization, the latter linked to subtle amino acid differences that dramatically affect biological distribution and function (Eyers et al., 2005, Bayliss et al., 2004, Fu et al., 2009). In vertebrates, the Aurora family of protein kinases are grouped into two functionally distinct sub-families composed of Aurora A and Aurora B and C (Carmena et al., 2009). Aurora kinases contain a conserved C-terminal kinase domain, which phosphorylates substrates contained within a minimal R-X-S/T consensus motif (Sardon et al., 2010, Haydon et al., 2003). The Aurora kinase domain is itself regulated by a critical autophosphorylation event in the activation loop (Ohashi et al., 2006, Eyers et al., 2003, Bayliss et al., 2003, Dodson et al., 2013), which is associated with catalytic activation and timely execution of different phases of mitosis (Hegarat et al., 2011, Tyler et al., 2007, Scutt et al., 2009, Yasui et al., 2004) and meiosis (Bury et al., 2017). During the cell cycle, Aurora A has canonical roles in centrosome duplication, spindle bipolarity, chromosome segregation and spindle checkpoint maintenance (Courtheoux et al., 2018), which together contribute to fundamental processes such as centrosomal and genomic stability and likely reflect the long-recognised oncogenic properties associated with overexpressed Aurora A (Bischoff et al., 1998). Interestingly, Aurora A has also recently been implicated in the control of energy production in cancer cells through mitochondiral targeting (Bertolin et al., 2018). In contrast, Aurora B is a centromeric kinase that is required for the spindle assembly checkpoint and rate-limiting for cytokinesis in cells, although it is also associated with cell transformation and a target for anti-proliferative therapy (Girdler et al., 2006). Aurora C is most highly expressed in germ cells, where it can functionally replace Auora B as a chromosome passenger protein (Carmena and Earnshaw, 2003).

Aurora A biology is controlled spatially *via* targeted subcellular localization mediated through the formation of dynamic protein complexes with non-catalytic binding partners such as TPX2 (Gruss et al., 2002, Bayliss et al., 2003, Eyers et al., 2003, Kufer et al., 2002, Wittmann et al., 2000), TACC3 (Burgess et al., 2018) and NMYC (Richards et al., 2016), which control distinct Aurora A signaling outputs relevant to cellular dynamics. Indeed, Aurora A/TPX2 holoenzyme complexes co-localise at the polar end of spindle microtubules, where Aurora A is maintained in dynamic pools alongside the mitoitc kinesin Eg5, which is localised by a C-terminal region of TPX2 (Eckerdt et al., 2008, Zeng et al., 2010, Kufer et al., 2002). Allosteric Aurora A activation on the spindle is thought to occur by binding to microtubule-associated proteins (which are also Aurora A substrates) such as TPX2 and TACC3 (Dodson and Bayliss, 2012). Mechanistically, Aurora A activation requires autophosphorylation of Thr 288 (and possibly Thr 287), which drives the kinase into the most catalytically-competent conformation (Walter et al., 2000, Eyers et al., 2003, Bayliss et al., 2004). Interestingly, the TPX2 complex also protects Aurora A from Thr 288 dephosphorylation by PP1 and PP2A phosphatases *in vitro* (Eyers et al., 2003, Bayliss et al., 2004), although accumulating evidence suggests that the PP6 Ser/Thr phosphatase is rate-limiting for mitotic dephosphorylation at this site in cells (Zeng et al., 2010). In addition to TPX2, Aurora A is targeted to core protein co-factors, including the centrosomal protein Cep192, the mitotic entry factor BORA, the ciliary protein Pifo and the transcription factor NMYC (Joukov and De Nicolo, 2018), whose interaction with a specific inactive Aurora A conformation is important for controlling NMYC stability in cells, and the basis for new approaches to target Aurora A output with drugs (Richards et al., 2017).

Structural analysis of Aurora A confirms the importance of activation loop dynamics for activity and binding-partner interactions (Burgess et al., 2018). Phosphorylation, as well as TPX2 binding, stabilise the activation segement in an appropriate conformation for catalysis, and distinct Aurora A conformations can also be induced and/or stabilised by a huge variety of chemical small molecules (Dodson et al., 2010, McIntyre et al., 2017, Pitsawong et al., 2018, Tyler et al., 2007, Scutt et al., 2009, Damodaran et al., 2017). Such compounds have been extremely useful to validate dozens of cellular Aurora A substrates (Tyler et al., 2005; Sardon, 2010), which include other cell cycle-regulated kinases such as PLK1 (Macurek et al., 2008, Scutt et al., 2009). Aurora A is also regulated by distinct post-translational modifications, including acetylation. For example, the Ubiquitin ligase CHFR ubiquitinates Aurora A *in vitro* and *in vivo*, targeting it for proteosome-mediated degradation(Yu et al., 2005). Interestingly, Aurora A is acetylated by arrest defective protein 1 (ARD1) acetyltransferase at lysines 75 and 125, and ARD1-mediated acetylation promotes cell proliferation and migration (Vo et al., 2017).

In the clinical context, overexpression of Aurora A is frequently detected in various human malignancies, including leukemia, breast, prostate, and colon cancers (Keen and Taylor, 2004, Bischoff et al., 1998, Zhou et al., 1998), and lower overall survival in patients with colorectal cancer correlates with increased levels of Aurora A (Goos et al., 2013). For these reasons, Aurora A has been pursued for several decades as a target for the development of novel anti-cancer therapeutic agents, some of which have show potential in the clinic. Beginning with MLN8054 (Manfredi et al., 2007), a number of potent, and highly selective, ATP-competitive small molecule inhibitors of Aurora A have been reported, with several advancing to phase II clinical trials, and MLN8237 (alisertib) (Sloane et al., 2010) proceeding to phase III evaluation after demonstrating activity in a variety of tumour settings (Tayyar et al., 2017, D’Assoro et al., 2015). Interestingly, while numerous ATP-competitive and allosteric ligands for Aurora A have been developed, some of which can specifically target the highly similar Aurora A and Aurora B, no type IV (covalent) inhibitors of Aurora kinases have yet been reported, despite clinical successes with covalent tyrosine kinase inhibitors such as ibrutinib, afatinib and osimertinib.

Coenzyme A (CoA) is essential for the viability of all living cells where it is a major regulator of cellular metabolism (Leonardi et al., 2005, Theodoulou et al., 2014). The CoA biosynthetic pathway is highly conserved in prokaryotes and eukaryotes and requires five enzymatic steps, involving sequential conjugation of pantothenic acid (vitamin B5), cysteine and ATP (Martinez et al., 2014, Sibon et al., 2016). The presence of a nucleotide moiety and a highly reactive thiol group permits broad diversity in biochemical reactions, which CoA employs to activate carbonyl-containing molecules and to generate covalent thioester derivatives such as Acetyl CoA, Malonyl CoA and 3-hydroxy-3-methylglutaryl CoA. Intracellular levels of CoA and its thioester derivatives are tightly regulated by a variety of stimuli, including hormones, nutrients, metabolites and stresses (Leonardi et al., 2005, Tsuchiya et al., 2014). The main rate-limiting enzyme in CoA biosynthesis is pantothenate kinase which initiates the biosynthetic pathway (Rock et al., 2000, Zhang et al., 2005, Dansie et al., 2014). Furthermore, the activity of the last enzyme in the pathway, CoA synthase, is regulated by extracellular stimuli and stress response (Zhyvoloup et al., 2003, Breus et al., 2009, Gudkova et al., 2012, Breus et al., 2010).

CoA and its thioester derivatives are strategically positioned at the crossroads of cellular anabolic and catabolic pathways, controling the Krebs cycle, ketogenesis, the biosynthesis of proteins, lipids and cholesterol, oxidation of fatty acids and degradation of amino acids. In additon, they are also involved in the regulation of gene expression and cellular metabolism via post-translational modifications, such as protein acetylation, butyrylation, malonylation and succinylation. Abnormal biosynthesis and homeostasis of CoA and its derivatives is associated with various human pathologies, including neurodegeneration, cancer, metabolic disorders and cardiac hypertrophy (McAllister et al., 1988, Reibel et al., 1981, Zhou et al., 2001, Dusi et al., 2014).

Protein CoAlation is a novel non-canonical function of CoA that involves covalent modification of cysteine residues in cellular response to oxidative and metabolic stress (Gout, 2018). This recent discovery was only possible with the development of unique research tools and methodologies, which revealed protein CoAlation as a widespread and reversible PTM involved in cellular redox regulation (Malanchuk et al., 2015). To date, more than one thousand proteins have been found to be CoAlated in a variety of prokaryotic and eukaryotic cells (Tsuchiya et al., 2017, Tsuchiya et al., 2018). Protein CoAlation occurs at a low level in exponentialy growing cells, but is strongly induced in reponse to various oxidising agents, including H_2_O_2_, diamide, menadione and t-butyl hydroperoxide. Protein CoAlation was shown to alter the molecular mass, charge, and activity of modified proteins, and to protect them from irreversible sulfhydryl overoxidation. These novel findings suggest that under normal growth conditions CoA mainly functions to produce metabolically active thioesters, while it may act as a low molecular weight antioxidant protecting protein cysteine thiols from irreversible overoxidation (Gout, 2018).

The presence of exposed cysteine residues on the surface of proteins provides a simple, often reversible, mechanism for post-translational modification of cell signaling proteins in response to a wide variety of redox conditions. Most protein classes contain surface-exposed Cys residues, and for the protein kinase superfamily, this knowledge has been exploited to successfully develop covalent (usually irreversible) ATP-dependent kinase inhibitors with clinical utility (Lanning et al., 2014; Liu Q et al., 2010). Interestingly, oxidative stress has marked effects on the cell cycle and proliferation (Chiu and Dawes, 2012), and redox regulation of signaling-active components such as tyrosine phosphatases (Tonks, 2005) and cell cycle phosphatases such as CDC25 (Savitsky and Finkel, 2002) contribute to redox-regulated checkpoints. More recently, direct effects of oxidative stress on abnormal spindle dynamics have been correlated with Aurora A inhibition (Wang et al., 2017). However, whether CoA regulates cell cycle-regulated protein kinases currently remains unknown.

In this study, we report direct Aurora A regulation through covalent interaction with CoA. We found that CoA is a specific ATP-competitive Aurora A inhibitor *in vitro*, and exploited a combination of biochemical, structural, mass spectrometry and cellular studies to confirm a preferential mode of CoA binding to the active kinase conformation. X-ray validation of the CoA binding mode confirms that it exploits Thr 217 in the ATP binding site, a locus that distinguishes Aurora A from Aurora B (Dodson et al., 2010, Sloane et al., 2010), to maintain a unique interaction with the 3’-phosphoADP moiety of CoA, positioning it for the unprecedented formation of a covalent bond between the pantetheine thiol and Cys 290, a remote residue found in the Aurora kinase activation loop. In support of this model, covalent modification of Aurora A by CoA was induced during cellular responses to oxidative stress, and microinjection of CoA perturbed spindle formation and chromosome alignment *in vivo*, causing potent cleavage arrest in fertilized mouse embryos. Taken together, our data reveal Aurora A as a novel target of the key metabolic regulator CoA, and open the door for the development of a new class of small molecules that exploits a ‘dual-anchor’ mechanism of inhibition bridging the Aurora A ATP-binding site and activation segment.

## Results

### Kinase profiling screen with coenzyme A reveals specific inhibition of Aurora A

CoA and ATP share an ADP moiety, which we hypothesised might lead to a serendipitous modulation of the activity of protein kinases by CoA. We tested this notion in a human kinome-wide activity-based screen (MRC National Centre for Protein Kinase Profiling, University of Dundee). Profiling of 117 protein kinases with CoA, dpCoA and ADP provided clear evidence that CoA is a relatively selective inhibitor for human Aurora A (Fig. 1A and Table 1). Moreover, CoA inhibited Aurora A with marked selectivity (87% Aurora A inhibition at 100 µM, n=3) when compared to ADP, a very non-specific kinase inhibitor that operates based on its close similarity to ATP. Only a few other kinases, including the tyrosine kinases SRC and YES, were inhibited ≤50% by CoA under these conditions. Interestingly, inhibition of Aurora B by CoA was also detected (46% inhibition at 100 µM compound), while Aurora C was not present in the screen (Table 1). Dephospho-CoA (dpCoA), which lacks the 3′-phosphorylated nucleotide, exhibited intermediate specificity between CoA and ADP, raising the possibility that both the pantetheine and 3′-phosphate moieties of CoA could be important for optimal inhibition of Aurora A (Fig. 1A). Further analysis confirmed that CoA inhibits active T288 phosphorylated (recombinant) Aurora A with an IC_50_ of approximately 5 µM in the presence of 5 M competing ATP *in vitro* (Fig. 1B). Using CoA Sepharose as an affinity matrix, we confirmed that endogenous Aurora A present in exponentially growing HepG2 cells interacts specifically with conjugated CoA, when compared to beads alone (Supplementary Fig. S1). Furthermore, bound Aurora A was efficiently eluted from CoA Sepharose with an excess of CoA.

**Figure 1.**
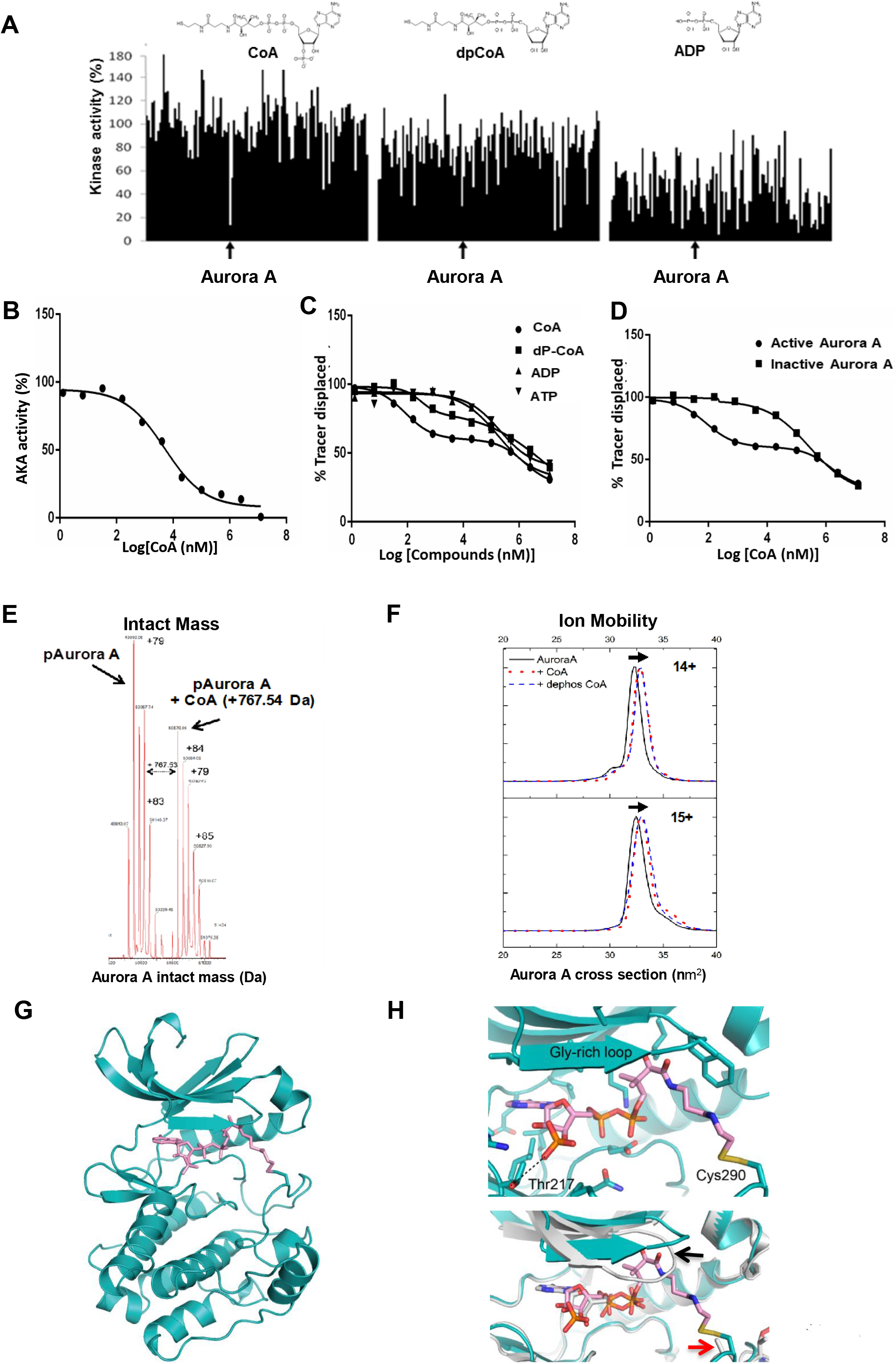
Coenzyme A binds directly to Aurora A and inhibits catalytic activity. **(A)** Kinase profiling screen reveals selective inhibition of Aurora A by CoA. The effect of CoA, dpCoA and ADP on the activity of 117 kinases was assayed using a radioactive filter-binding assay at the International Centre for Kinase Profiling, Dundee University. Each compound was tested at 100 μM final concentration in the presence of indicated concentrations of ATP for each kinase (Table 1). Schematic structures of the tested compounds are shown above. **(B)** CoA inhibits Aurora A *in vitro*. Kinase activity of recombinant full-length His-Aurora A was assayed by measuring incorporation of γ^33^P-ATP into myelin basic protein in the presence of 5 µM ATP and an 11-point serial dilution of CoA. **(C)** Analysis of binding kinetics of CoA, dpCoA, ADP and ATP towards active (pT288-phosphorylated) Aurora A using a Lanthascreen Eu Kinase Binding assay **(D)** CoA preferentially binds to the active, pT288 phosphorylated form of Aurora A, when compared to the catalytically inactive dephosphorylated kinase. Dephosphorylation of His-Aurora A was carried out in the presence of recombinant PP1 phosphatase. **(E)** Intact mass analysis of phosphorylated Aurora A incubated with CoA in the absence of DTT. Covalent incubation of CoA with Aurora A generates a population of phosphorylated Aurora A (pAurora A) alongside covalent adducts containing an extra mass attributable to CoA (pAurora A + CoA). **(F)** Ion Mobility spectra and calculated cross sectional area (^TW^CCS_N2→He_) for [M+14H]^14+^ and [M+15H]^15+^ ions of pT288 Aurora A measured in the absence (black line) or presence of CoA (red lines) or dephospho-CoA (blue lines). Overlapping conformations of Aurora A are shown, the more extended of which have an increased mean cross sectional area associated with the presence of CoA and dephospho-CoA. **(G)** Crystal structure of Aurora A (teal) in complex with CoA (pink). **(H)** Upper panel, magnified view of CoA and its interactions with Aurora A, highlighting the side-chains of Thr 217 and Cys 290, forming a covalent bond with CoA. Lower panel, superposition of Aurora A/CoA with Aurora A/ADP (PDB code 1OL7), showing the shifted position of the Gly-rich loop (black arrow) and the equivalent position of Cys 290 (red arrow). Note that the side chain of Cys 290 was modelled without the sulfur atom in the Aurora A/ADP structure, due to weak electron density.

**Table 1.** Kinase profiling of CoA, dpCoA and ADP. The ability of CoA, dpCoA or ADP (100 μM final concentration of each compound and the indicated concentrations of ATP) to inhibit the phosphotransferase activity of 117 kinases was determined using a radioactive filter-binding assay at the International Centre for Kinase Profiling, DSTT, University of Dundee. Kinase activity in the absence of inhibitor is taken as 100%.

| Kinase | CoA | ADP | dpCoA | ATP in assay (μM) |
| --- | --- | --- | --- | --- |
| ABL | 111.2 | 22.9 | 98.0 | 5 |
| AMPK | 90.8 | 54.1 | 75.9 | 50 |
| ASK1 | 129.1 | 70.2 | 88.6 | 50 |
| Aurora A | 13.4 | 38.0 | 29.9 | 5 |
| Aurora B | 54.3 | 47.8 | 55.1 | 20 |
| BRK | 114.1 | 41.3 | 83.1 | 20 |
| BRSK1 | 110.4 | 42.8 | 96.4 | 20 |
| BRSK2 | 114.7 | 52.4 | 88.8 | 50 |
| BTK | 67.9 | 18.2 | 30.6 | 50 |
| CAMK1 | 123.6 | 80.1 | 125.6 | 50 |
| CAMKKα | 97.4 | 52.9 | 71.4 | 20 |
| CDK2-Cyclin A | 93.2 | 26.3 | 81.1 | 20 |
| CHK1 | 122.6 | 47.6 | 78.8 | 20 |
| CHK2 | 88.9 | 43.2 | 56.0 | 20 |
| CK1δ | 80.8 | 57.6 | 80.4 | 20 |
| CK2α | 80.1 | 5.4 | 39.8 | 5 |
| CLK2 | 108.9 | 33.9 | 80.1 | 5 |
| CSK | 92.6 | 18.5 | 54.9 | 20 |
| DAPK1 | 106.2 | 10.3 | 64.4 | 5 |
| DYRK1A | 93.1 | 75.9 | 90.2 | 50 |
| DYRK2 | 102.4 | 42.4 | 77.0 | 50 |
| DYRK3 | 95.1 | 16.2 | 91.2 | 5 |
| EF2K | 67.1 | 86.2 | 103.0 | 5 |
| EPH-A2 | 102.0 | 45.0 | 99.2 | 50 |
| EPH-A4 | 114.6 | 32.5 | 75.6 | 50 |
| EPH-B1 | 116.9 | 11.6 | 80.0 | 20 |
| EPH-B2 | 117.3 | 14.3 | 91.1 | 20 |
| EPH-B3 | 104.2 | 18.0 | 94.4 | 20 |
| EPH-B4 | 106.8 | 9.8 | 62.1 | 50 |
| ERK1 | 118.2 | 67.7 | 114.0 | 5 |
| ERK2 | 119.7 | 54.4 | 81.7 | 20 |
| ERK8 | 101.4 | 16.6 | 38.5 | 5 |
| FGF-R1 | 60.0 | 41.7 | 31.4 | 20 |
| GCK | 108.8 | 56.5 | 71.2 | 20 |
| GSK3β | 94.7 | 18.2 | 72.1 | 5 |
| HER4 | 100.5 | 29.1 | 75.7 | 5 |
| HIPK1 | 115.3 | 31.7 | 92.8 | 20 |
| HIPK2 | 153.3 | 16.5 | 73.4 | 5 |
| HIPK3 | 123.1 | 34.2 | 91.5 | 20 |
| IKKβ | 90.5 | 26.6 | 48.0 | 5 |
| IKKε | 125.2 | 73.8 | 91.4 | 50 |
| IRAK4 | 89.1 | 82.1 | 74.8 | 20 |
| JAK2 | 105.5 | 7.1 | 62.5 | 5 |
| JNK1 | 85.4 | 53.9 | 82.4 | 20 |
| JNK2 | 93.9 | 57.3 | 76.1 | 20 |
| JNK3 | 101.3 | 58.6 | 63.2 | 20 |
| Lck | 100.4 | 35.1 | 69.0 | 50 |
| LKB1 | 103.3 | 71.0 | 81.1 | 20 |
| MAPKAP-K2 | 93.1 | 53.0 | 81.7 | 20 |
| MAPKAP-K3 | 98.4 | 25.7 | 63.0 | 20 |
| MARK1 | 95.3 | 28.1 | 68.0 | 20 |
| MARK2 | 93.1 | 25.0 | 81.5 | 20 |
| MARK3 | 88.9 | 14.8 | 46.2 | 5 |
| MARK4 | 87.7 | 49.1 | 86.6 | 50 |
| MEKK1 | 96.3 | 68.1 | 105.8 | 20 |
| MELK | 76.3 | 63.5 | 104.0 | 50 |
| MINK1 | 106.3 | 49.4 | 103.9 | 50 |
| MKK1 | 107.2 | 16.4 | 52.3 | 5 |
| MKK2 | 103.0 | 35.4 | 53.0 | 5 |
| MKK6 | 88.6 | 39.1 | 88.2 | 50 |
| MLK1 | 109.9 | 41.4 | 76.0 | 20 |
| MLK3 | 70.2 | 46.8 | 61.9 | 20 |
| MNK1 | 150.3 | 60.7 | 84.2 | 50 |
| MNK2 | 137.5 | 41.2 | 87.1 | 50 |
| MSPK1 | 101.8 | 94.1 | 101.7 | 50 |
| MSK1 | 106.4 | 53.3 | 70.5 | 20 |
| MST2 | 118.8 | 40.5 | 72.5 | 20 |
| MST4 | 107.0 | 56.9 | 89.7 | 20 |
| NEK2α | 148.5 | 42.1 | 100.4 | 50 |
| NEK6 | 118.4 | 26.6 | 69.6 | 50 |
| NUAK1 | 98.3 | 32.4 | 68.3 | 20 |
| p38α MAPK | 112.5 | 60.3 | 87.7 | 50 |
| p38β MAPK | 159.6 | 51.1 | 94.5 | 20 |
| p38δ MAPK | 128.0 | 37.3 | 71.3 | 5 |
| p38γ MAPK | 127.2 | 26.0 | 65.9 | 5 |
| PAK2 | 88.8 | 90.3 | 111.9 | 20 |
| PAK4 | 69.9 | 73.8 | 75.9 | 5 |
| PAK5 | 70.6 | 79.5 | 86.0 | 20 |
| PAK6 | 98.4 | 78.4 | 100.7 | 20 |
| PDK1 | 113.3 | 23.3 | 91.2 | 20 |
| PHKγ1 | 105.1 | 75.6 | 90.9 | 50 |
| PIM1 | 103.0 | 46.1 | 97.7 | 20 |
| PIM2 | 111.0 | 9.8 | 62.9 | 5 |
| PIM3 | 121.7 | 36.1 | 96.7 | 20 |
| PKA | 98.9 | 27.8 | 81.3 | 20 |
| PKCα | 104.2 | 38.6 | 76.8 | 5 |
| PKCβ | 118.3 | 79.2 | 105.1 | 50 |
| PKCα | 78.1 | 76.5 | 88.9 | 20 |
| PKCζ | 99.3 | 37.7 | 89.9 | 5 |
| PKCγ | 70.2 | 71.9 | 66.4 | 20 |
| PKD1 | 111.9 | 56.2 | 104.3 | 50 |
| PLK1 | 101.3 | 52.2 | 100.0 | 5 |
| PRAK | 99.7 | 15.2 | 77.7 | 20 |
| PRK2 | 71.3 | 20.8 | 67.5 | 5 |
| RIPK2 | 131.4 | 46.0 | 88.4 | 20 |
| ROCK 2 | 94.3 | 57.8 | 96.4 | 20 |
| RSK1 | 97.2 | 58.5 | 98.9 | 50 |
| RSK2 | 96.5 | 47.9 | 62.9 | 50 |
| S6K1 | 89.1 | 35.5 | 30.5 | 20 |
| SGK1 | 102.8 | 38.7 | 74.3 | 20 |
| SmMLCK | 92.1 | 73.0 | 91.3 | 20 |
| Src | 43.8 | 31.8 | 8.9 | 50 |
| SRPK1 | 116.3 | 60.9 | 89.1 | 50 |
| STK33 | 71.8 | 54.6 | 83.3 | 50 |
| SYK | 98.3 | 70.9 | 105.9 | 20 |
| TAK1 | 115.3 | 15.8 | 79.6 | 5 |
| TAO1 | 113.2 | 34.0 | 98.3 | 20 |
| TBK1 | 106.3 | 95.1 | 109.6 | 50 |
| TIE2 | 93.0 | 33.1 | 83.4 | 20 |
| TTK | 101.3 | 54.7 | 75.8 | 20 |
| YES1 | 49.0 | 18.6 | 11.5 | 20 |
| ZAP70 | 89.7 | 13.2 | 79.8 | 5 |
| IGF-1R | 93.0 | - | - | 5 |
| IR | 100.0 | - | - | 20 |
| IRR | 92.0 | - | - | 5 |
| TrkA | 104.0 | - | - | 20 |
| VEGFR | 81.0 | - | - | 20 |

Next, we quantified the binding kinetics of CoA, dpCoA, ADP and ATP for catalytically active (T288 phosphorylated) and catalytically inactive (non-phosphorylated) forms of Aurora A. A Lanthascreen Eu Kinase FRET Binding Assay was employed to evaluate the interaction of CoA with Aurora A, and tease apart any molecular contributions made by the pantetheine tail and nucleotide 3’-phosphate. We found that the displacement curves for CoA and dpCoA were biphasic, indicating the presence of two binding sites with higher and lower affinities towards CoA (Fig. 1C). Bacterially expressed Aurora-A proteins auto-phosphorylate themselves to a varying degree, resulting in a heterogeneous mixture containing active and inactive Aurora-A (Scutt et al., 2009). Importantly, treatment of the Aurora-A preparation with protein phosphatase 1, which results in T288 dephosphorylation (Eyers et al., 2003), caused disappearance of the high affinity-binding site, confirming that this was related to the activation (phosphorylation) state of the kinase (Fig. 1D). We next determined the affinity of interaction by titrating Aurora A with CoA, dpCoA and ADP, and compared them with values obtained from *in vitro* kinase (phosphotransferase) assays (Table 2). Under the assay conditions used, the apparent IC_50_ for the interaction of active Aurora-A with CoA and dpCoA were 82 nM and 294 nM, respectively, whilst the IC_50_ value for ADP was approximately 177 µM. Taken together, these results clearly demonstrate that CoA and dpCoA preferentially bind to the active (T288-phosphorylated) form of Aurora A, and suggest an involvement of the pantetheine tail and the the 3′-phopshate of CoA in mediating a potentially specific interaction with Aurora A.

**Table 2.** CoA preferentially binds to active, T288 phosphorylated Aurora A. The binding of T288 phosphorylated Aurora A (‘active’), or unphosphorylated Aurora A (‘inactive’), was assessed for each compound and the inhibitory constants were calculated. The data confirm that both the pantetheine and the 3’ phosphate of CoA participate in the interaction between Aurora A and CoA.

| Compound | Aurora A (active)<br>IC<sub>50</sub> (nM), binding<br>assay | Aurora A (inactive)<br>IC<sub>50</sub> (nM), binding<br>assay | Aurora A (active)<br>IC<sub>50</sub> (μM), activity<br>assay |
| --- | --- | --- | --- |
| ATP | 208 | 2990 | - |
| ADP | 177 | 1280 | 54 |
| dpCoA | ~0.294 | $7 \times 10^{17}$ | 17 |
| CoA | ~0.082 | 283 | 5 |

### Biophysical and structural analysis of the Aurora A:CoA complex

To confirm the interaction between CoA and Aurora A using intact mass spectrometry, we pre-incubated pT288 (phosphorylated) Aurora A with excess CoA, and observed a sub-stoichiometric shift for each of the differentially phosphorylated intact Aurora A species present in the preparation, consistent with the formation of phosphorylated Aurora A adducts, each containing a single molecule of CoA (Mr 767.54 Da, Fig. 1E). We also evaluated effects of CoA and dpCoA on the gas phase conformation of Aurora A using Ion Mobility Mass Spectrometry (IM-MS), which is a new approach for probing kinase conformational dynamics (Byrne et al., 2016). The calculated cross-sectional area of intact Aurora A (nm^2^) in both 14^+^ and 15^+^ charge states increased significantly after pre-incubation with both CoA or dpCoA, consistent with their ability to bind to Aurora A and induce a conformational change. Comparative IM-MS of each complex confirmed species with larger cross-sectional areas than the control, whose formation was independent of the 3’ phosphate group, predicting that Aurora A conformational changes are likely to be driven by covalent attachment of the CoA pantetheine tail (Fig. 1F).

To gain insights into the precise binding mode of the interaction between CoA and active Aurora A, we next determined the crystal structure of the complex to 2.5 Å resolution (Table 3, Fig. 1G). As predicted, CoA occupies the canonical ATP binding site of Aurora A, between the N-lobe and C-lobe of the kinase. Remarkably, the extended CoA pantetheine moiety stretches well away from the ATP site towards the kinase activation loop, forming a disulphide bond with the side chain of Cys 290 adjacent to the phosphorylated residue Thr 288 (Fig. 1H). Unexpectedly, the pantothenic acid moiety of CoA interacts with the tip of the Gly-rich loop (aa142-145) and displaces it from the position found in Aurora-A/ADP complexes, consistent with our IM-MS data. Interestingly, the 3’ phosphate group of CoA is also ideally positioned to form a H-bond with the side chain of Thr 217, which is evolutionary conserved in vertebrate Aurora A, but not Aurora B (Dodson et al., 2010, Sloane et al., 2010). Interestingly, our crystal structure of Aurora A/CoA is entirely consistent with independent molecular modelling of Aurora A bound to CoA, based on the previously determined structure of Aurora A/ADP (Supplementary Fig. S2). The structure employed for modelling (PDB code 1OL7), was chosen based on previous knowledge that the side chain of Cys 290 is in a potentially suitable position to react chemically with the pantetheine tail of CoA. Importantly, most features of the experimental crystal structure were predicted by this model, including the specific interaction of the 3’ phosphate group of CoA with the side chain of Thr 217. One exception is the experimentally up-shifted position of the Gly-rich loop in the presence of CoA, which was unexpected based on the model, where the Gly-rich loop resides in a canonical position, packed against the ADP 5’ phosphate groups.

**Table 3.** Data collection and refinement statistics.

| Aurora A 122-403/CoA |  |
| --- | --- |
| **Data collection** |  |
| Space group | *P*6<sub>1</sub>22 |
| Cell dimensions |  |
| *a*, *b*, *c* (Å) | 80.89, 80.89, 173.34 |
| $\alpha$, $\beta$, $\gamma$ (°) | 90.00, 90.00, 120.00 |
| Resolution range (Å) | 44.57-2.50 (2.54-2.50)* |
| *R*<sub>merge</sub> (%) | 8.58 (171) |
| *I*/σ*I* | 21.92 (1.61) |
| Completeness (%) | 100 (100) |
| Redundancy | 17.59 (15.07) |
| **Refinement** |  |
| Resolution (Å) | 44.57-2.50 |
| No. reflections | 12263 |
| *R*<sub>work</sub> / *R*<sub>free</sub> | 20.44/25.21 |
| No. atoms |  |
| Protein | 2102 |
| Hetero | 73 |
| Water | 26 |
| Mean *B*-factors |  |
| Protein | 69.90 |
| Hetero | 77.39 |
| Water | 69.30 |
| Wilson *B*-factor | 70.56 |
| r.m.s. deviations |  |
| bond lengths (Å) | 0.013 |
| bond angles (°) | 1.238 |
| **MolProbity analysis** |  |
| All-atom clash-score | 9.17 |
| Rotamers outliers (%) | 0 |
| Ramachandran outliers (%) | 0 |
| Ramachandran favoured (%) | 95.06 |
| MolProbity score | 1.83 |

\*Values in parentheses are for highest-resolution shell.

### Aurora A is CoAlated on Cys 290 *in vitro*

The electron density map unambiguously supports the formation of a covalent bond between the sulfur atoms in the side chain of Cys290 and pantetheine thiol of CoA (Fig. 2A). To investigate the contribution of this bond to the interaction, we employed a validated *in vitro* CoAlation assay (Tsuchiya et al., 2017). Here, recombinant Aurora A was pre-incubated with CoA, and the reaction mixture was separated by SDS-PAGE under non-reducing conditions followed by immunoblotting with a monoclonal anti-CoA antibody. As shown in Fig. 2B, CoA interacts with Aurora A in a DTT-sensitive manner, suggesting a thiol-dependent covalent mode of binding. Ponceau staining of immunoblotted samples revealed that the mobility of Aurora A was retarded when CoA disulphide was present in the reaction mixtures. The addition of DTT completely abolished the observed electrophoretic mobility shift, indicating the formation of CoA–Aurora A mixed disulphides. LC-MS/MS analysis was employed to map CoA-modified cysteine(s) of *in vitro* CoAlated Aurora A. This analysis revealed that Cys290 is CoAlated in the RTpTLC^290^GTLDYPPEMIEGR peptide (Supplementary Fig. S3A), confirming the co-existence of Thr 288-phosphorylated (active) Aurora A and Cys 290-CoA *in vitro*. We also examined whether binding of the 3′-phosphate ADP moiety of CoA to the Aurora A ATP binding pocket primes the pantetheine thiol for disulphide bond formation with Cys290 in the activation loop. *In vitro* CoAlation of Aurora A was carried out with 100 µM CoA and increasing concentrations of ATP (0-10 mM). Aurora A CoAlation gradually decreased in the presence of increasing concentrations of ATP, and was completely abolished by inclusion of 1 mM DTT (Fig. 2C). These observations suggest that occupancy of the ATP binding pocket by the 3’-phosphate ADP moiety of CoA is a pre-requisite for covalent modification of Aurora A (see below).

**Figure 2.**
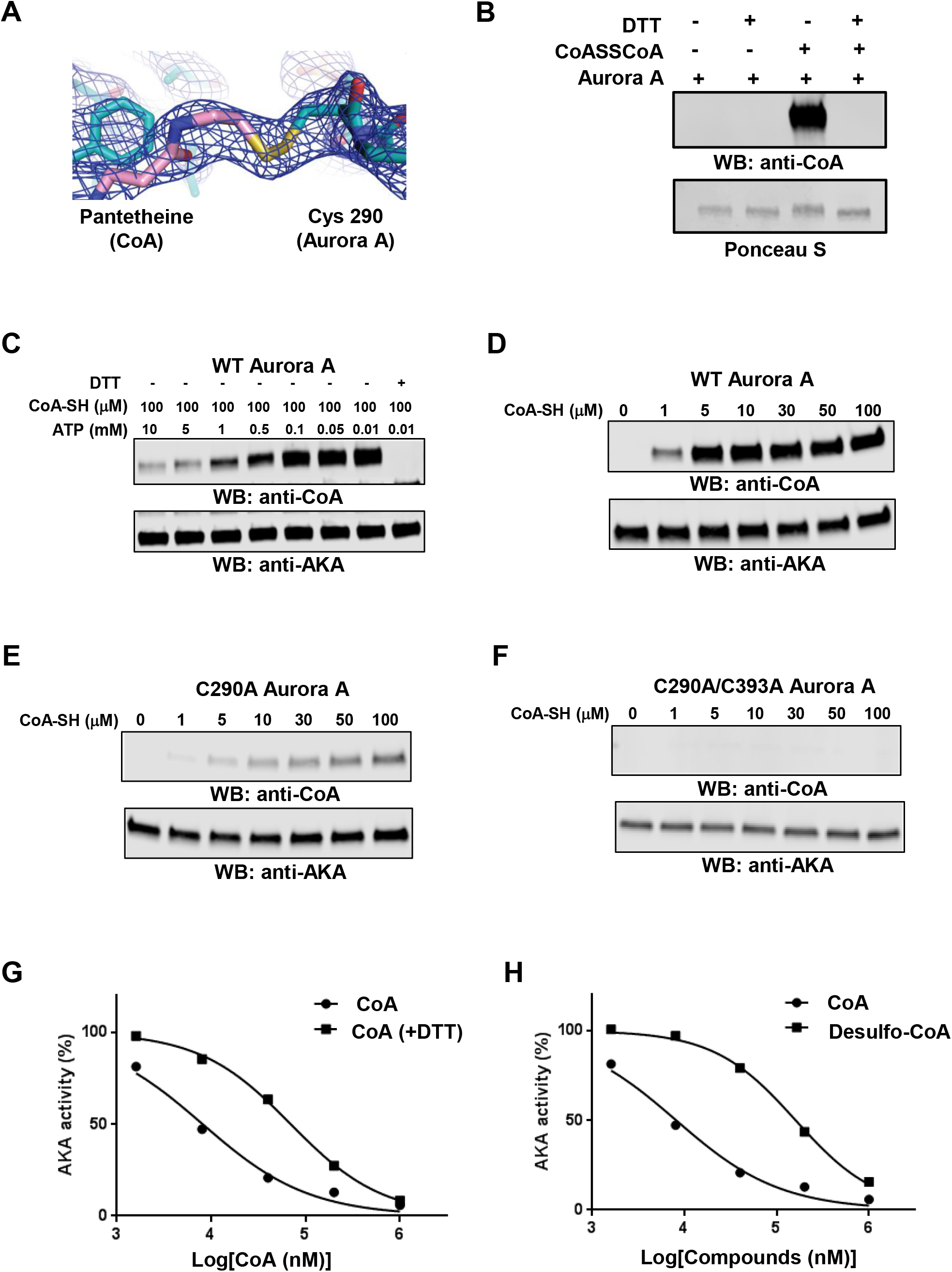
CoA reversibly binds to Aurora A in an ATP-competitive manner through Cys 290. **(A)** Electron density around the pantotheine tail region of CoA and Cys 290 of Aurora A, 2Fo-Fc map contoured at 1 σ. **(B)** Aurora A is covalently modified by CoA in a DTT-sensitive manner. *In vitro* CoAlated Aurora A was separated by SDS-PAGE in the presence or absence of DTT and immunoblotted with anti-CoA antibody. The membrane was stained with Ponceau red to visualize total Aurora A. **(C)** Binding of CoA to the ATP-binding pocket is required to facilitate covalent modification of Aurora A by CoA. *In vitro* CoAlation of Aurora A was carried out in the presence of 100 µM CoA and the indicated concentration of ATP. Generated samples were separated by SDS-PAGE in the presence or absence of DTT and immunoblotted with anti-CoA and anti-Aurora A antibodies. **(D)** WT Aurora A is efficiently CoAlated by CoA *in vitro*. **(E)** The C290A Aurora A mutant exhibits significantly reduced binding of CoA when compared to WT Aurora A. **(F)** The C290/393A Aurora A mutant is not covalently modified by CoA in vitro. *In vitro* CoAlation of WT Aurora A, C290A and C290/393A mutants was performed with the indicated concentration of CoA. The reaction mixtures were separated by SDS-PAGE and immunoblotted with anti-CoA and Aurora A antibodies. **(G)** The inhibitory effect of CoA towards Aurora A is reduced by DTT. Recombinant His-Aurora A was used to determine the IC_50_ value for CoA in the presence or absence of 1 mM DTT. **(H)** Reduced inhibitory effect of desulfo-CoA compared to CoA. Desulfo-CoA lacks the reactive SH group at the end of the pantetheine tail.

Aurora A has two surface-exposed cysteine residues present in the kinase domain: Cys290 in the activation loop and Cys393 in the C-terminal regulatory region (Burgess and Bayliss, 2015). Mutational analysis allowed us to demonstrate that covalent modification of the Cys290Ala mutant by CoA is dramatically reduced in comparison to wild-type Aurora A, while no binding was detected with the double Cys290/393Ala mutant (Fig. 2D-F). The dose-dependency of inhibition of active Aurora A by CoA was also examined in the absence and presence of a reducing agent (Fig. 2G). Aurora A was inhibited by CoA under both experimental conditions. However, the IC_50_ value (CoA) was an order of magnitude higher in the presence of DTT (47 μM), when compared to that obtained in the absence of DTT (5 μM). To explore the mode of regulation further, we tested the inhibitory effect of desulpho-CoA on Aurora A kinase activity. In this analogue of CoA, the reactive SH group at the end of the pantetheine tail is absent and unable to covalently modify Aurora A. Consistently, the dose response inhibition curve for desulpho-CoA was also shifted, resulting in a significantly higher IC_50_ value for Aurora A (160 μM), confirming the involvement of the CoA thiol group in the specific inhibition of Aurora A kinase activity (Fig. 2H).

### The role of Thr 217 in mediating a selective interaction between Aurora A and CoA

We next sought to identify the basis of selectivity of CoA for Aurora A over other kinases. To do this, we focussed on the closely related Aurora A and B kinases, which share 70% sequence identity in the kinase domain (Fig. 3A). The putative hydrogen bond between the -OH side chain of Thr 217 in Aurora A and the 3′-phosphate of the ribose ring of CoA (Fig. 1H) was of particular interest, since Thr 217 is known to contribute to the specificity of MLN8054, a selective Aurora A inhibitor (Sloane et al., 2010), where the corresponding residue Glu 177 in Aurora B prevents compound binding, likely due to electrostatic repulsion (Sloane et al., 2010, Dodson et al., 2010). We compared covalent modification of Aurora A and Aurora B using an *in vitro* CoAlation assay (Fig. 3B, D). This analysis revealed strong CoAlation of Aurora A, while Aurora B was CoAlated at an extremely low level under optimal Aurora A assay conditions (100 µM CoA). Mutational analysis showed that the Thr217Glu mutant of Aurora A exhibits significantly reduced CoA binding compared to the wild-type kinase (Fig. 3C). In contrast, Glu177Thr Aurora B mutant exhibited strong binding to CoA (Fig. 3E). Together, these data indicate that both the 3′-phosphate ADP moiety and the thiol group of CoA are involved in mediating specificity-determining interactions with Aurora A *via* targeting of Thr 217 and Cys290 respectively.

**Figure 3.**
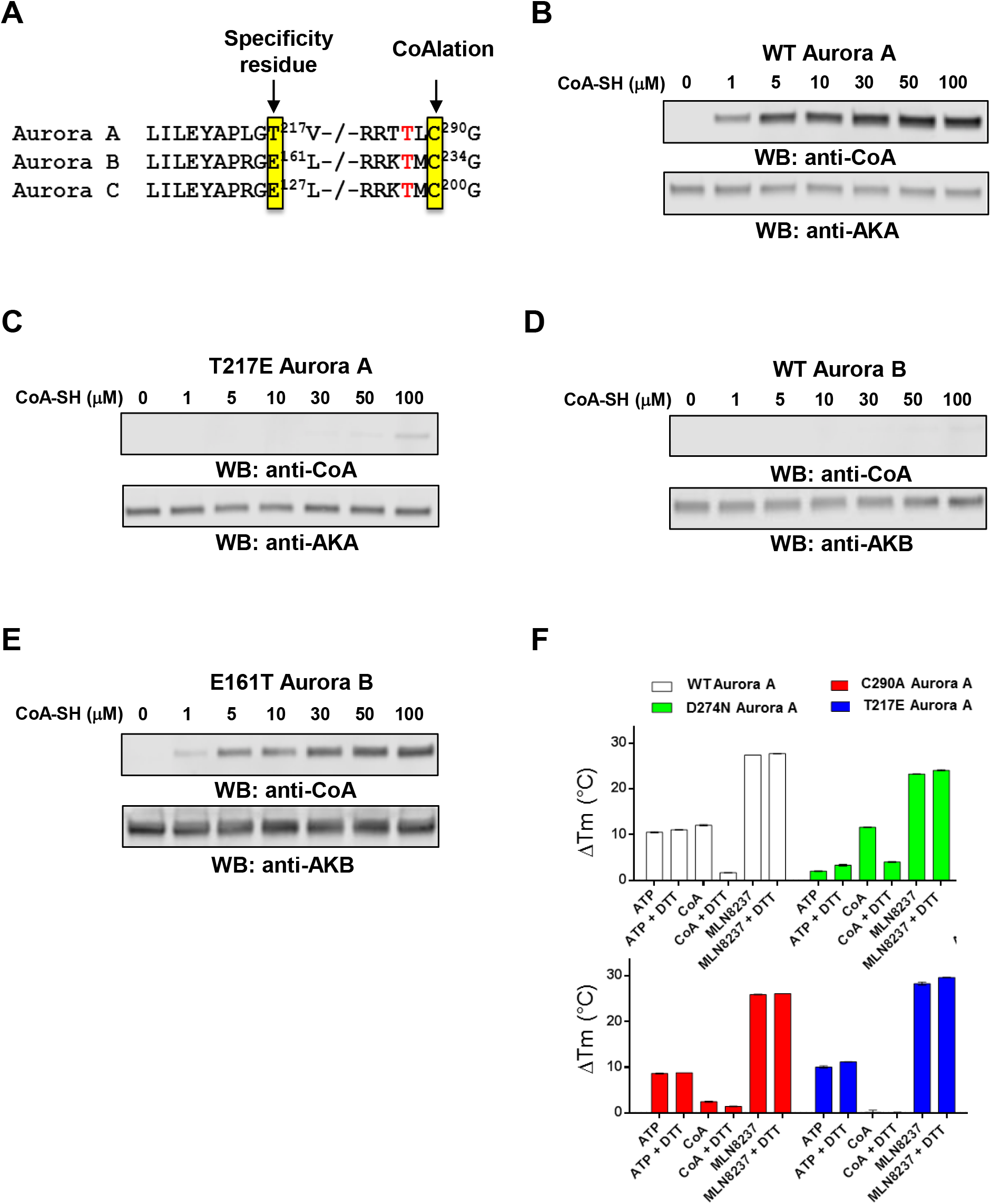
Specificity of Aurora A interaction with CoA is controlled by Thr 217. **(A)** Amino acid conservation in vertebrate Aurora kinases. Thr 217 defines Aurora A, and is changed to a Glu residue in Aurora B and C. Cys 290 (boxed) is invariant in all Aurora kinases, and lies in the activation segment adjacent to the phosphorylated Thr 288 (human Aurora A numbering). **(B)** *In vitro* CoAlation of Aurora A. **(C)** *In vitro* CoAlation of Aurora A is abolished in the Thr217Glu mutant. **(D)** *In vitro* CoAlation is not detected with Aurora B. **(E)** The Glu161Thr Aurora B mutant is efficiently CoAlated. Experiments were performed with the indicated concentration of CoA. Reaction mixtures were separated by SDS-PAGE and immunoblotted with anti-CoA, Aurora A or Aurora B antibodies.**(F)** A thermal shift assay was employed to evaluate Aurora A binding to 5 mM ATP, 5 mM CoA or 0.1 mM MLN8237, in the presence and absence of 1 mM DTT, as indicated. Assays were performed with T288 phosphorylated, active, Aurora A (open bars), kinase-dead, dephosphorylated Aurora A (Asp274Asn, green bars), Cys290Ala Aurora A (red bars) or Thr217Glu Aurora A (blue bars). Mean ΔT_m_ values ± SD (n=3) were calculated by subtracting the control T_m_ value (buffer, no addition) from the measured T_m_ value.

### Thermal shift analysis of the Aurora A/CoA interaction

To probe thermal effects of CoA in the presence and absence of DTT, we profiled ligand binding in solution (Fig. 3F). WT Aurora A is phosphorylated and catalytically active, whereas D274N Aurora A cannot bind to ATP and therefore remains dephosphorylated after isolation from bacteria (Haydon et al., 2003). C290A Aurora A lacks the thiol-containing Cys residue in the kinase activation segment (Fig. 3A), whereas T217E Aurora A mimics Aurora B by swopping a polar Thr for a charged Glu (see above). Thermal melting profiles (Foulkes et al., 2018) for each protein were obtained in the presence and absence of DTT and ATP, CoA or the Aurora A inhibitor MLN8237. Binding of ATP, CoA and MLN8237 induced a marked change in thermal stability of WT Aurora A, which was dependent on non-reducing conditions in the case of CoA, and entirely consistent with findings with Aurora A isolated from human cells (Fig. 2B). Interestingly, CoA was still able to interact with kinase-dead D274N (inactive) Aurora A in a DTT-dependent manner, although its interaction with Mg-ATP was severely blunted, as expected (green bars). In contrast, C290A Aurora A bound to ATP, but was deficient in CoA-induced thermal stabilization (red bars). Finally, a CoA-induced thermal shift was completely abolished in the T217E Aurora A mutant, in contrast to those of either ATP or MLN8237, reiterating the importance of a Thr at this position for CoA binding.

### Binding of TPX2 protects Aurora A from covalent modification and inhibition by CoA

In vertebrates, TPX2 enhances Aurora A autophosphorylation and protects it from dephosphorylation by phosphatases. Biochemical and crystallographic analyses have revealed the molecular mechanism by which the N-terminal fragment of TPX2 (residues 1–43) interacts with Aurora A, and locks it in a stable active conformation, also protecting it from dephosphorylation by PP1 phosphatase (Fig. 4A, left). Protection from enzymatic dephosphorylation (Eyers et al., 2003) occurs because the phosphate group of Thr 288 is buried in the TPX2-bound form (Bayliss et al., 2003). The side chain of Cys290 is also buried in the TPX2-bound, active conformation of Aurora A, and furthermore it points away from the ATP binding side, whereas the thiol group is exposed in structures of active Aurora A obtained in the absence of TPX2 in which CoA (Fig. 4A, middle) or ADP (Fig. 4A, right) are bound. We therefore investigated whether the inhibitory effect of CoA on Aurora A was modulated by TPX2. Dose inhibition curves of Aurora A by CoA were generated in the presence and absence of TPX2. As shown in Fig. 4B, the presence of the TPX2 1-43 fragment in an *in vitro* kinase assay completely abrogated the inhibitory effect of CoA on Aurora A. However, pre-incubation of Aurora A with CoA prevented TPX2-induced kinase activation in a DTT-sensitive manner (Fig. 4C). These findings indicate that binding of TPX2 protects Aurora A not only from dephosphorylation of Thr 288 by phosphatases, but that it can also prevent covalent modification (and therefore inhibition) of the kinase by CoA.

**Figure 4.**
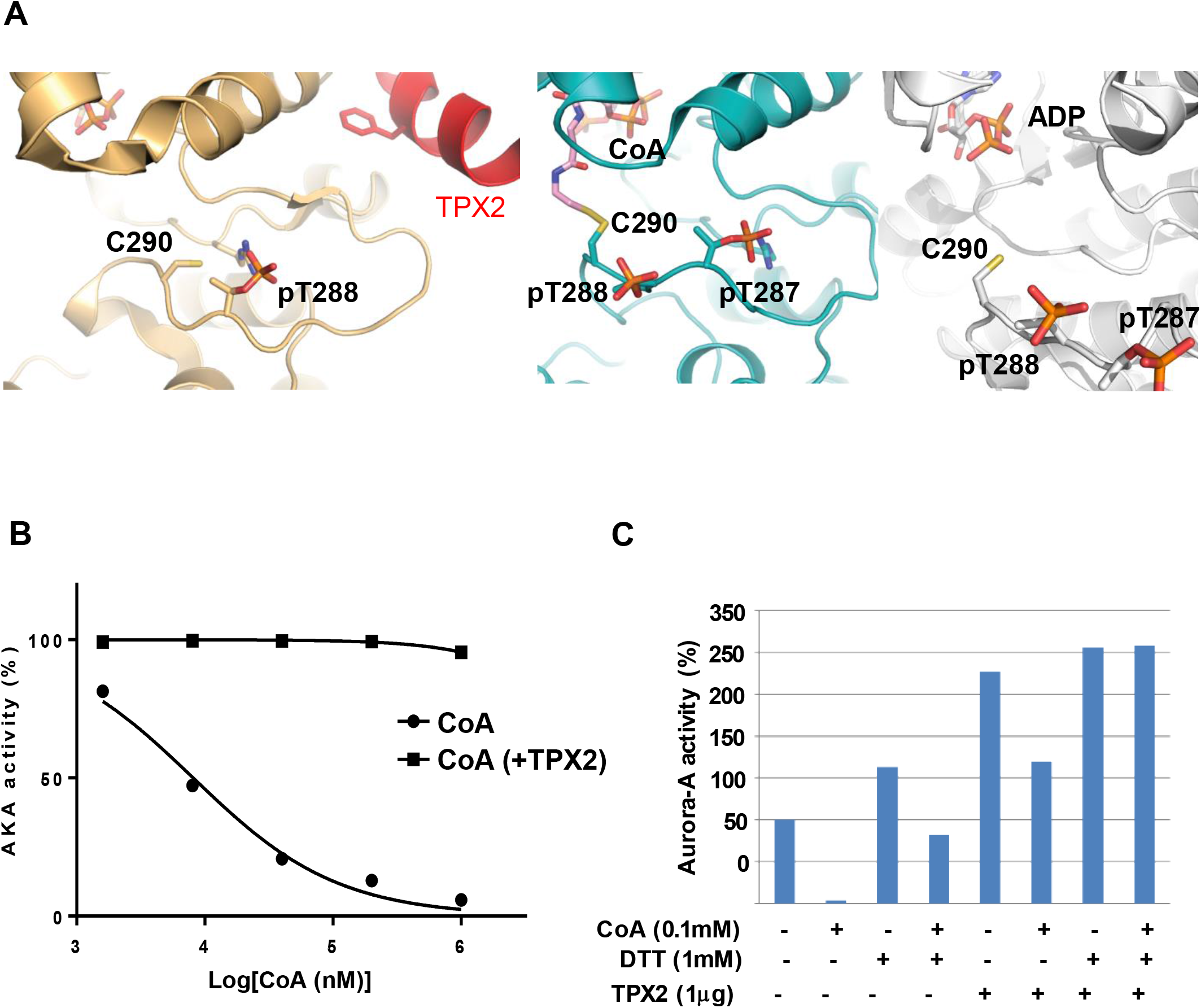
Inhibitory effect of TPX2 binding on CoA interaction with Aurora A and position of pT288. **(A)** Magnified views of the activation loop of Thr 288-phosphorylated Aurora A in the crystal structures of complexes with: ADP/TPX2, PDB code 1OL5; CoA (center); or ADP, PDB code 1OL7 (right). Note that in the ADP complex, the sulfur on the side chain of Cys290 is not included in the experimental model but is shown here, and the β3-αC loop removed for clarity. **(B)** Aurora A is resistant to inhibition by CoA in the presence of TPX2 1-43 peptide. Kinase activity of recombinant His-Aurora A was assayed by measuring incorporation of γ^33^P-ATP into myelin basic protein in the presence or absence of N-terminal fragment of TPX2 (residues 1–43) and serial dilutions of CoA **(C)** Pre-incubation of Aurora A with CoA inhibits TPX2-induced kinase activation in a DTT-sensitive manner.Recombinant His-Aurora A was preincubated with 100 M CoA before the *in vitro* kinase assay which was performed in the presence or absence of the N-terminal fragment of TPX2 (residues 1–43) and DTT.

### Oxidative stress induces Aurora A CoAlation in cells

Extensive protein CoAlation is observed in mammalian cells and tissues exposed to oxidative and metabolic stress (Tsuchiya et al., 2017). We therefore examined whether covalent modification of Aurora A by CoA is induced in cells after exposure to various oxidative stresses (Tsuchiya et al., 2017). We exploited HEK293/Pank1β cells, which stably overexpress pantothenate kinase 1 β (Pank1β), the major rate-limiting enzyme in CoA biosynthesis (Tsuchiya et al., 2017). It has been shown previously that Pank1β overexpression induces a significant increase in CoA biosynthesis. These cells are also ideal for evaluating protein CoAlation in cellular response to oxidative and metabolic stress (Tsuchiya et al., 2017). Accordingly, cells were transiently transfected with plasmids encoding N-terminally FLAG-tagged Aurora A and Aurora B, and then treated with 500 μM H_2_O_2_ prior to lysis. Immunoprecipitation of transiently expressed FLAG-tagged proteins followed by immunoblotting with anti-CoA antibody revealed specific CoAlation of Aurora A in cells after exposure to H_2_O_2_, but Aurora B was poorly CoAlated (Fig. 5A). To map the site(s) of Aurora A CoAlation, transiently overexpressed FLAG-Aurora A was immunoprecipitated from cells exposed to H_2_O_2_, processed and analysed by LC-MS/MS. This analysis revealed a CoAlated Aurora A peptide derived from the activation segment with the sequence TTLC^290^GTLDYLPPEMIRGR, confirming covalent modification in cells (Supplementary Fig. S3B), and consistent with *in vitro* CoAlation of recombinant Aurora A (Fig. 2B). Next, Aurora A CoAlation was examined in cells exposed to a diverse range of oxidising agents. As shown in Fig. 5B, CoAlation of transiently expressed Aurora A was strongly induced in cells treated with diamide and to a lesser extent after exposure to menadione or phenylarsine oxide (PAO). It was recently reported that ROS-induced cell cycle arrest promotes hyperphosphorylation of Aurora A leading to abnormal mitotic spindle assembly and significant mitotic delay (Wang et al., 2017). Therefore, we examined phosphorylation of Aurora A at Thr 288 by probing the same immunoprecipitation samples with anti-pT288 antibody. We confirmed that treatment of cells with H_2_O_2_ and menadione promoted an increase in Aurora A phosphorylation at Thr 288. We also observed a strong induction of phosphorylation with PAO, when compared to control cells (Fig. 5B). Taken together, these results allow us to propose a simple model for regulation of Aurora A by oxidative stress in cells (Fig. 5C). In exponentially growing cells, Aurora A carries out many of its cellular functions by using ATP as substrate for protein phosphorylation. The binding of CoA to the Aurora A ATP binding pocket via the ADP moiety is effectively outcompeted by ATP (Fig. 5C, left panel). In contrast, exposure of cells to oxidative stress induces covalent binding of CoA to phosphorylated Aurora A (Fig. 5C, right panel), which is accompanied by a loss of catalytic activity and altered cell signaling.

**Figure 5.**
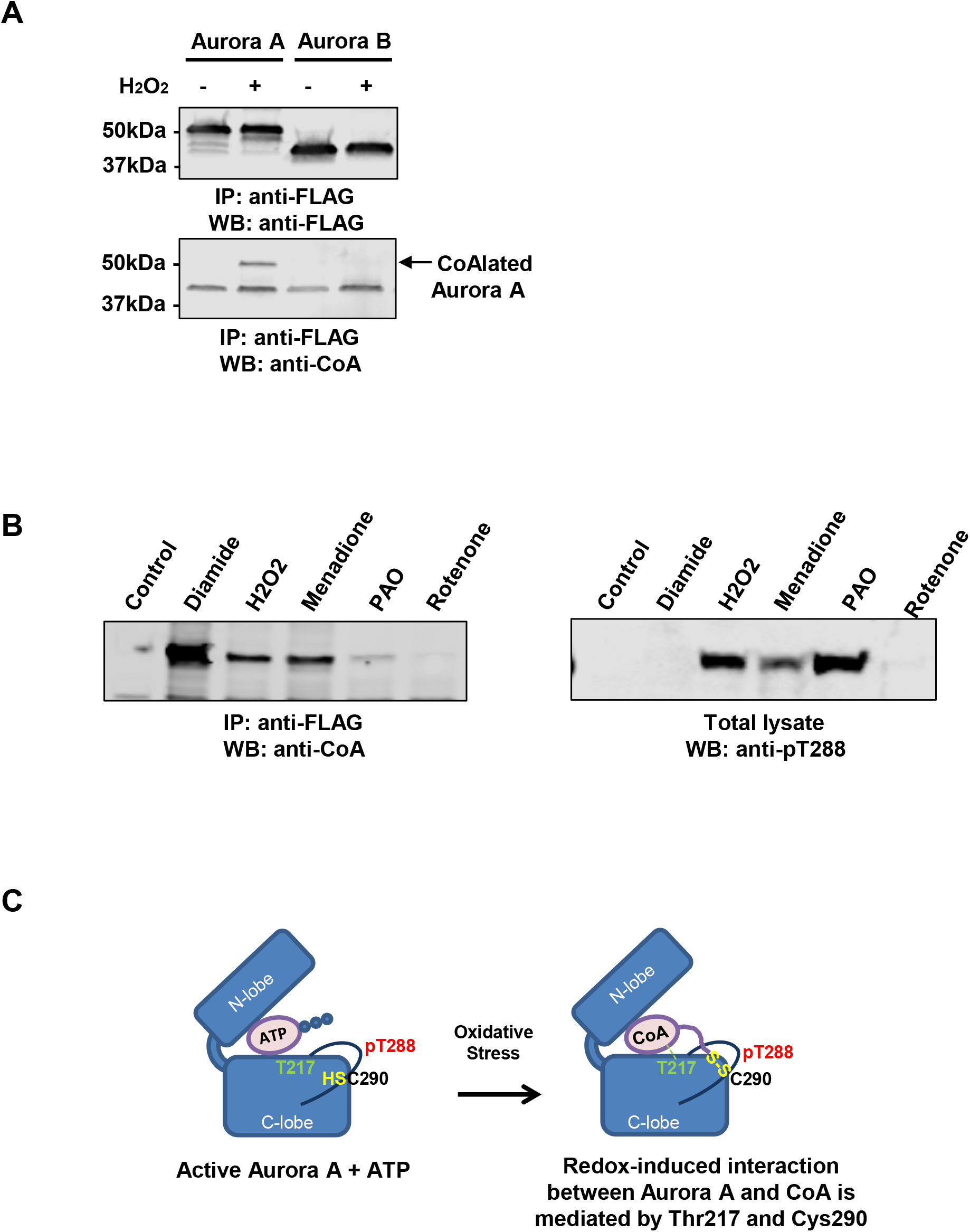
Oxidative stress induces Aurora A CoAlation in human cells. **(A)** Aurora A CoAlation is induced in cellular response to H_2_O_2_. FLAG-tagged WT Aurora A and WT Aurora B were transiently overexpressed in HEK293/Pank1β. Transfected cells were treated for 30 min with 0.25 mM H_2_O_2_. Overexpressed proteins were immunoprecipitated with an anti-FLAG antibody and the immune complexes immunoblotted with anti-CoA and anti-FLAG antibodies. **(B)** Oxidising agents promote Aurora A CoAlation and phosphorylation at Thr 288 *in vivo*. FLAG-tagged WT Aurora A was transiently overexpressed in HEK293/Pank1β. Transfected cells were treated for 30 min with a panel of oxidising agents (250 μM H_2_O_2_, 500 μM diamide, 50 μM menadione, 10 μM phenylarsine oxide and 100 nM rotenone). Transiently expressed proteins were immunoprecipitated with an anti-FLAG antibody, separated by SDS-PAGE under non-reducing conditions and immunoblotted with anti-CoA and anti-pT288 Aurora A antibodies. **(C)** Schematic illustration showing the key features of the ‘dual anchor’ mechanism for interaction of CoA with Thr 217 and Cys 290 in Aurora A

### Microinjection studies in mouse oocytes

It has been established that functional (catalytically-active) Aurora A is essential for normal spindle formation during mouse oocyte maturation (Bury et al., 2017, Saskova et al., 2008). To investigate the effect of CoA on Aurora A functions *in vivo*, we employed a well-established model for microinjecting CoA and related molecules into immature murine oocytes. Oocytes arrested at the GV stage were injected with different concentrations of CoA. 17 hours after GV release, oocytes were fixed and stained for α-tubulin (green) and DNA (blue) to observe effects on spindle formation. We found that microinjection of CoA markedly increased the number of oocytes with atypical spindles and misaligned chromosomes in a dose-dependent manner (Supplementary Fig. 4A). Our *in vitro* studies demonstrated the importance of both the pantetheine tail and the 3’ phosphate of CoA in mediating a high-affinity interaction with Aurora A (Fig. 1). We therefore examined the effect of injected CoA or different compounds representing integral parts of the CoA molecule to cause abnormal spindle formation and misalignment of chromosomes in mouse oocytes. We found that CoA, but not pantetheine, ADP or AMP, was capable of disrupting bipolar spindle formation and chromosome alignment *in vivo* (Supplementary Fig. 4B). These data confirm that the pantetheine tail is a requirement for the inhibitory effects of CoA in mouse oocytes. Taken together, our *in vivo* oocyte microinjection studies provide strong evidence that an increase in cellular CoA has the potential to deregulate spindle formation and the alignment of chromosomes in oocytes, causing meiotic cleavage arrest.

## Discussion

Binding partners and post-translational modifications regulate the activity of Aurora A by switching it between inactive and active conformations (Levinson, 2018). The intrinsic catalytic activity of Aurora A is low prior to mitosis, and activation involves autophosphorylation of Thr 288 in the activation loop, which is promoted through interaction with microtubule-associated proteins such as TPX2 and TACC3. In this study, CoA is identified as a key metabolic molecule that binds and inhibits Aurora A activity, potentially as part of the cellular response to oxidative stress. Aurora A CoAlation was not observed in exponentially growing cells, but it was significantly induced as part of the cellular response to oxidative stresses. Based upon our ‘dual-anchor’ inhibitory model (Fig. 5C), Aurora A CoAlation is initiated by anchoring the ADP moiety of CoA to the ATP-binding pocket of the oxidized form of Aurora A, which involves a selectivity-determining interaction between the 3′-phosphate ADP moiety of CoA and Thr 217. Wedging of the ADP moiety then positions the flexible pantetheine tail of CoA with the thiol group in close proximity for covalent bond formation with Cys 290 in the Aurora A activation segment, two residues C-terminal from the regulatory phosphorylated Thr 288 residue. We speculate that under oxidative stress, covalent binding of the pantetheine thiol to Cys 290 keeps the ADP moiety firmly anchored to the ATP binding pocket, making it inaccessible to ATP, and preventing phosphotransferase activity. The “dual anchor” mechanism of locking the kinase in an inactive state therefore makes CoA a more potent and selective endogenous inhibitor of Aurora A when compared to Aurora B and other protein kinases.

### What is the biological function of Aurora A CoAlation?

In addition to an inhibitory effect on Aurora A activity, CoAlation at Cys 290 may serve other regulatory purposes. First, CoA modification might protect Cys 290 from overoxidation, which may lead to an irreversible loss of function and subsequent inactivation and/or degradation of Aurora A. The widespread ability of CoA to act as a low-molecular-weight antioxidant in response to oxidative and metabolic stress in prokaryotic and eukaryotic cells has been recently demonstrated (Tsuchiya et al., 2017, Tsuchiya et al., 2018). In these studies, *in vitro* CoAlation of the catalytic Cys 151 in glyceraldehyde-3-phosphate dehydrogenase (GAPDH) was shown to protect this essential glycolytic enzyme against irreversible overoxidation and the associated loss of activity. It remains to be seen whether oxidation of Cys 290 in Aurora A to sulphenic acid (which is reversible) facilitates covalent modification by CoA via a disulphide-exchange mechanism, protecting this highly susceptible thiol from irreversible overoxidation. Secondly, covalent modification of Cys 290 by CoA may modulate the phosphorylation status of Thr 288, which is critical for Aurora A catalytic activity. Previous studies reported differential sensitivity of Thr 288 to dephosphorylation by phosphatase complexes of the phosphoprotein phosphatase (PPP) family. An intriguing possibility is that redox-induced covalent linkage of CoA to Cys 290, which is directly adjacent to Thr 288, may prevent the binding of phosphatases that control the dephosphorylation of Thr 288. In support of this hypothesis, we found that phosphorylation of Aurora A at Thr 288 is significantly increased under oxidative stress induced by H_2_O_2_, menadione and PAO. However, it is also possible that a build-up of phosphorylated Aurora A is due to the fact that CoA is a more efficient inhibitor of the phosphorylated form of Aurora A, meaning that it spares Aurora A autophosphorylation, or that CoAlation of Aurora A changes recognition of the protein by the T288 phosphospecific antibody.

An involvement of CoA, a fundamental metabolic integrator in the regulation of Aurora A, provides new insight into our understanding of regulatory mechanisms of cell cycle progression mediated by metabolic adaptation to oxidative stress. Energy flow and biosynthetic processes are tightly regulated during the cell cycle. CoA plays a critical role in generating a diverse range of metabolically active thioester derivatives (including Acetyl-CoA, HMG-CoA, Malonyl-CoA and Acyl-CoA), which function as driving forces for generating ATP, the biosynthesis of macromolecules and the regulation of gene expression *via* protein acetylation. The presence of a highly reactive thiol group in CoA and its potential involvement in redox regulation has intrigued researchers for many decades, but progress in this field of study has been hampered by a lack of precise research tools and methodologies. Our findings are consistent with novel functions of CoA in the cellular response to oxidative and metabolic stress, which involves covalent attachment of this coenzyme to redox-sensitive Cys residues (Tsuchiya et al., 2017). CoA levels also change during early embryogenesis in *Xenopus laevis*, consistent with shifts in energy usage and perhaps CoAlation before and after the midblastula transition (Tsuchiya et al., 2014). Protein CoAlation is therefore emerging as an important post-translational modification implicated in redox regulation of a diverse range of proteins (Gout, 2018), including redox-controlled protein kinases (Corcoran and Cotter, 2013).

Our work also demonstrates a hitherto unappreciated role for CoA in the regulation of Aurora A activity in cellular response to oxidative stress, and also suggests the potential importance of this form of regulation for other protein kinases that possess cysteine residues within the activation segment, or elsewhere (Zhao et al., 2017). This includes the Aurora A-related basophilic Ser/Thr kinase PKA, for which inhibition by CoA was not detected *in vitro*. However, the reversible glutathionylation of Cys 199, equivalent to Cys 290 in Aurora A, has previously been demonstrated (Humphries et al., 2002) and correlated with functional roles for the Cys 199 redox status in controlling T-loop phosphorylation at the adjacent Thr 197 site in PKA, equivalent to Thr 288 in Aurora A (Humphries et al., 2005, Humphries et al., 2007). In addition to Aurora A, CoA also modestly inhibits several unrelated protein kinases, including SRC, YES, BTK and FGFR1 (Table 1). However, none of these tyrosine kinases contains a Cys residue in the activation segment, and it remains to be seen whether covalent modification of these kinases by CoA occurs in cells, and if so, whether it is also induced by oxidative stress. In the case of CoA, our data suggest that the “dual anchor” inhibitory mechanism in Aurora A permits precise spatial communication between the ATP-binding site and the activation loop, explaining the very high degree of specificity for CoAlation of Aurora A and preventing high-affinity interaction with even closely-related kinases, such as Aurora B.

### Significance

Our findings raise the interesting possibility that new covalent approaches can be developed to target Aurora A, or other kinases possessing Cys residues in the activation segment, with small inhibitory molecules. This dual targeting mechanism might be the basis of a new approach for design of specific, and potentially irreversible, Aurora A small molecule inhibitors, with long-time target engagement in cells. The majority of protein kinase (including Aurora A) inhibitors are currently ATP competitive (non-covalent) inhibitors, which are designed to gain at-least some selectivity through the recognition of unique features of conformation-specific ATP-binding pockets. However, there has been a resurgence of interest in generating covalent compounds targeting Cys residues in drug targets, most notably RAS (Ostrem et al., 2013), previously considered undruggable. Moreover, biochemical tool compounds (Zhang et al., 2016, Ocasio et al., 2018) and clinical inhibitors covalently targeting oncogenic kinase, are enjoying significant clinical success (Zhao and Bourne, 2018). However, one of the liabilities of such compounds is the appearance of drug resistance, caused by point mutations in the Cys residue at the covalent drug interface. In this study, we report a new ‘dual-mode’ of Aurora A inhibition by the metabolic integrator CoA, which also reveals how specific targeting of kinases might be accomplished with a distinct class of irreversible inhibitor, whose ability to covalently trap kinases in an inhibitory conformation could represent a new way of tuning signaling outputs.

## Acknowledgments

We thank the members of Cell Regulation laboratory at the Department of Structural and Molecular Biology (UCL) for their valuable inputs throughout this study; UCL Darwin Research Facility for tissue culture and analytical biochemistry support. We thank Dr Matthias Vonderach and Dr Philip Brownridge for help with MS and Sam Evans for outstanding technical support. We thank the support staff of Diamond beamline I03. L.G. thanks University of Hyderabad for research facilities. This work was funded by grants to I.G. (UCLB 13-014 and 11-018; BBSRC BB/L010410/1); P.E, (North West Cancer Research CR1088/1097) and R.B. (Cancer Research UK C24461/A23302); V.F. (National Academy of Sciences of Ukraine 0110U000692).

## Author Contributions

This study was conceived by I.G.; Y.T., A.Z., J.B., Y.H., T.T., F.B. and D.P.B. performed cell biology experiments, biochemical and biophysical assays; S.G.B. and R.B. designed and carrier out crystallization, structure determination and analysis of the Aurora-A/CoA complex; S.F. performed ion mobility and intact mass spectrometry; E.C. and L.G. carried out molecular docking studies; O.G. and V.F. developed and characterised anti-CoA Mabs; J.B. carried out mouse oocytes microinjection experiments; S.Y.P-C. and M.S. designed and performed the MS-MS experiments; Y.T., P.E., R.B., C.E.E., J.C., A.T. and I.G. designed analysed experiments; I.G., P.E. and R.B. wrote the manuscript with the assistance and approval of all authors.

## Declaration of Interests

The Authors declare that there are no competing interests associated with the manuscript.

## Materials and Methods

### Reagents and chemicals

The generation and characterization of the monoclonal anti-CoA antibody 1F10 has been described previously (Tsuchiya et al., 2017). All common chemicals and biochemicals were obtained from Sigma-Aldrich unless otherwise stated, including CoA, dpCoA, dsCoA, ATP, and ADP. The following antibodies were employed: mouse anti-CoA antibody; mouse anti-FLAG M2 antibody (Sigma-Aldrich); rabbit anti-Aurora A and anti-Aurora B antibodies (Merck-Millipore); rabbit anti-pT288 Aurora A (Cell Signaling Technology), Alexa Fluor 680 goat anti-mouse IgG H&L (Life Technologies) and IRdye 800 CW goat anti-rabbit IgG H&L (LI-COR Biosciences).

### Kinase profiling screen and Aurora A IC_50_ determination

Profiling of 117 protein kinases with CoA, dpCoA and ADP was carried out at the International Centre for Protein Kinase Profiling at the University of Dundee. Each compound was tested *in vitro*, in duplicate, at 100 μM final concentration using recombinant kinases, model substrates and optimal concentrations of ATP. Bacterially expressed full-length His-Aurora A was used to determine the IC_50_ value for CoA (in the presence or absence of DTT), dpCoA, ADP and desulfo-CoA. The assay was performed in the presence of 5 μM competing ATP. To examine whether the inhibitory effect of CoA on Aurora A is modulated by TPX2, IC_50_ values for CoA were also generated in the presence and absence of 1 μg TPX2 (residues 1-43) fragment.

### *In vitro* Aurora A kinase assay

Aurora A kinase activity was assayed by measuring incorporation of γ^33^P-ATP into myelin basic protein (Sigma). Purified recombinant protein (100 ng) was incubated at room temperature for 30 min in a total volume of 15 μl containing 50 mM HEPES pH 7.5, 10 mM MgCl2, 1mM EGTA, 0.05% Brij-35, 0.5 mg/ml myelin basic protein, and 5 μM γ^33^P-ATP (100-10,000 dpm/pmol). The reaction was stopped by spotting the reaction mixture onto squares of P81 phosphocellulose ion exchange paper (Whatmann), which were then immersed in 1.5% (v/v) phosphoric acid. After washing twice in 1 % phosphoric acid followed by two washes in distilled water, the papers were air dried and radioactivity was counted by a scintillation counter and recorded by Quantasmart version 2.03.

### *In vitro* CoAlation Assay

For CoAlation assays, 0.5 µg of purified recombinant preparations of wild type and mutant forms of His-Aurora A (Cys290Ala; Cys393Ala, Cys290/393Ala, Thr217Glu) and His-Aurora B (Glu161Thr) were incubated with a mixture of oxidised and reduced forms of CoA (CoASH and CoASSCoA, 1 mM final) in 20 mM Tris-HCl, pH 7.5 for 30 min at RT. The reaction was stopped by adding SDS gel-loading buffer without DTT, but containing 10 mM NEM. To examine whether binding of the 3′-phosphate ADP moiety of CoA to the Aurora A ATP binding-pocket co-ordinates the pantetheine thiol for disulphide bond formation with Cys290, the *in vitro* CoAlation assay was carried out with 100 µM CoA and increasing concentrations of ATP (0-10 mM).

### CoA Sepharose pull-down assay

Exponentially growing HepG2 cells were lysed on ice in buffer, containing 50 mM Tris-HCl pH 7.5, 150 mM NaCl, 5 mM ethylenediaminetetraacetic acid (EDTA), 50 mM sodium fluoride (NaF), 5 mM tetra-sodium pyrophosphate (Na_4_P_2_O_7_) and 1% (v/v) Triton X-100, supplemented with 1x Protease Inhibitor Cocktail (PIC, Roche). After centrifugation, the supernatant was incubated on a rotating wheel with CoA Sepharose or Sepharose alone for 2 hr at 4^°^C. Beads were then washed extensively with lysis buffer and bound proteins were eluted from beads with 2x SDS loading buffer. To elute proteins that bound specifically to CoA Sepharose, the cell lysis buffer was supplemented with 100 μM CoA (CoA eluted fraction). Eluted proteins were separated by SDS-PAGE and Western blotted with anti-Aurora A antibody.

### Lanthascreen Eu Kinase FRET Binding Assay

A Lanthascreen Eu Kinase FRET Binding Assay (Invitrogen) was employed to determine IC_50_ values for the interaction of Aurora A with CoA, dpCoA, ATP, and ADP. In this assay, we used bacterially expressed His-Aurora A in active and inactive (PP1-treated, pT288 dephosphorylated) states. Recombinant Aurora A was incubated with Europium-conjugated anti-histidine tag antibody and an Alexa Fluor 647-labelled tracer, which binds to the ATP binding pocket of Aurora A. The close proximity of the anti-histidine (epitope-tag) antibody and the tracer results in a high degree of FRET (fluorescence resonance energy transfer) from the europium donor fluorophore to the Alexa Fluor 647 acceptor fluorophore. ATP-competitive inhibitors, such as CoA, displace the tracer from the active site, causing a loss of FRET signal. Assay set-up was performed as described by the manufacturer. Briefly, the time-resolved fluorescence resonance energy transfer assay (TR-FRET) was performed in black, low volume 384 well plates (Corning). Each well contained 5 nM kinase, 2 nM Eu-anti-His antibody and 10 nM kinase tracer 236 in kinase buffer A (50 mM Hepes pH 7.5, 10 mM MgCl2, 1 mM EDTA, 0.01% (v/v) Brij-35), varying amounts of CoA, dpCoA, ATP, and ADP. The reaction mix was incubated for 1 h at room temperature. The signal was measured at 665/620 nm emission ratio using BMG LABTECH plate reader (PHERAstar). All assays were performed using duplicates. The 11-point response curves were generated using GraphPad Prism software from the inhibition data generated.

### Plasmids and Protein purification for Differential Scanning Fluorimetry (DSF)

For enzyme and DSF assays, full-length human Aurora A and Aurora B and 1-43 TPX2 were produced in BL21 (DE3) pLysS *E. coli* cells (Novagen) with expression induced with 0.5 mM IPTG for 18 h at 18°C and purified as *N*-terminal His6-tag fusion proteins by immobilized metal affinity chromatography (IMAC) and size exclusion chromatography using a HiLoad 16/600 Superdex 200 column (GE Healthcare) equilibrated in 50 mM Tris/HCl, pH 7.4, 100 mM NaCl, 10 % (v/v) glycerol and 1 mM DTT. Asp274Asn and Thr217Glu Aurora A mutants were generated by PCR-site directed mutagenesis, expressed and purified as described previously (Sloane et al., 2010). Cys290Ala Aurora A was generated using standard procedures, and purified as described.

### Differential Scanning Fluorimetry

Thermal-shift assays were performed with a StepOnePlus Real-Time PCR machine (Life Technologies) using Sypro-Orange dye (Invitrogen) and thermal ramping (0.3 °C in step intervals between 25 and 94°C). All proteins were diluted to a final concentration of 5 μM in 50 mM Tris/HCl, pH 7.4 and 100 mM NaCl in the presence or absence of the indicated concentrations of ligand (final DMSO concentration no higher than 4 % v/v). CoA, dephosphoCoA or kinase inhibitors diluted from 10 mM DMSO stocks (Byrne et al., 2018) were assayed as described previously (Foulkes et al., 2018). Normalized data were processed using the Boltzmann equation to generate sigmoidal denaturation curves, and average T_m_/∆T_m_ values calculated as previously described (Murphy et al., 2014) using GraphPad Prism software.

### Molecular docking

Docking of CoA in the crystal structure of Aurora A kinase (PDB_ID: 1OL7) (Bayliss et al., 2003) was carried out using CDOCKER (Discovery Studio 3.1, Accelrys Inc). In the crystal structure, a 7Å grid was selected around the ATP binding site in order to define the location of probable CoA binding site. Twenty-five conformations of CoA were generated and ten docking poses were analysed.

### Crystallization and structure determination of the Aurora A 122-403/CoA complex

Aurora A (amino acids 122-403) was produced as described in earlier work (Burgess et al., 2015). The kinase was subject to size exclusion chromatography on a HiLoad 16/600 Superdex 200 pg column (GE Healthcare) equilibrated in 20 mM Tris pH 7.0, 0.2 M NaCl, 5 mM MgCl_2_, 10% (v/v) glycerol prior to crystallization trials. Aurora A was concentrated to 16.5 mg/ml and incubated with 5 mM CoA for 30 minutes at 30 °C. Crystallization screens were laid down in 96-well MRC plates using a Mosquito LCP crystallization robot (TTP Labtech) and incubated at 295 K. Crystals were produced in 0.03 M MgCl_2_.6H_2_0, 0.03 M CaCl_2_.2H_2_0, 0.1 M MOPS/HEPES-Na pH 7.5, 20% (v/v) PEG 500 MME, 10% (w/v) PEG 20 000 (Molecular Dimensions) and flash frozen directly from the drop.

X-ray diffraction data were collected on beamline I03 at Diamond Light Source, Oxford, England from a single crystal. Data processing was performed by the *xia2 3dii* automated data-reduction platform at Diamond (Winter et al., 2013). The structure of the complex was solved by molecular replacement using Phaser-MR (McCoy et al., 2007) and the structure of Aurora A 122-403 C290A, C393A (PDB 4CEG) as a model (Burgess and Bayliss, 2015). Phenix.refine was used to solve the structure and carry out iterative refinement (Adams et al., 2002). Model building was performed using Coot (Emsley and Cowtan, 2004). Structure validation and data quality were determined by Molprobity (Chen et al., 2010).

### Human cell culture

HEK293 and HEK293/Pank1β cells were cultured in Dulbecco’s Modified Eagle Medium (DMEM) (Lonza) supplemented with 10% foetal bovine serum (FBS) (Hyclone), 50 U/ml penicillin and 0.25 µg/ml streptomycin (Lonza). HepG2 cells were cultured in Williams E media (Lonza) supplemented with 10% FBS, 50 U/ml penicillin and 0.25 µg/ml streptomycin. All cell lines were tested and shown to be free of mycoplasma infection. Generation of HEK293/Pank1β cell line with stable overexpression of Pantothenate kinase 1β was previously reported (Tsuchiya et al., 2017).

### Transient transfection of HEK293/PANK1β cells and treatment with oxidizing agents

HEK293/Pank1β cells were transfected at ~60 % confluence with pCDNA3.1/FLAG-Aurora A and pCDNA3.1/FLAG-Aurora B plasmids, according to the manufacturer’s protocol using Turbofect reagent (Thermo Scientific). Transfected cells were allowed to grow for 24 h in complete DMEM with 10% FBS. The medium was replaced with pyruvate-free DMEM supplemented with 5 mM glucose and 10% FBS and cells were incubated for another 24 h. Pyruvate was removed from the media because it can act as an antioxidant and inactivate ROS. Cells were then treated with H2O2 (250 µM), menadione (50 µM), phenylarsine oxide (PAO, 10 µM), diamide (500 µM) or rotenone (1µM) for 30 min at 37 °C in pyruvate-free DMEM supplemented with 5 mM glucose. Cells were harvested by pressure washing and centrifuged at 1,800 g for 5 min at RT.

### Cell lysis, immunoprecipitation and Western blot analysis

Harvested cells were lysed at 4 °C for 20 min in lysis buffer, containing 50 mM Tris-HCl pH 7.5, 150 mM NaCl, 5 mM EDTA, 50 mM NaF, 5 mM Na4P2O7 and 1% Triton X-100, supplemented with fresh 100 mM NEM and fresh 1x PIC. Total cell lysates were centrifuged at 20817*x* g for 10 min at 4°C and the supernatant was collected for further analysis. Protein concentration was measured using the Bicinchoninic acid Protein Assay Kit (Thermo Scientific). Immunoprecipitation of transiently expressed FLAG-Aurora A and FLAG-Aurora B from cell lysates was carried out using Protein G Sepharose (Generon) and anti-FLAG antibody (Sigma-Aldrich). Proteins were eluted from beads with 2x SDS loading buffer and separated by SDS-PAGE under non-reducing conditions. Resolved proteins were transferred to a PVDF membrane (Bio-Rad Laboratories), which was then blocked with Odyssey blocking buffer. The membrane was incubated in anti-CoA or anti-FLAG antibodies for 2 h at room temperature (RT) or overnight at 4 °C, and then with secondary antibodies for 30 min (RT). Immunoreactive bands were visualised using Odyssey Scanner CLx and Image Studio Lite software (LI-COR Biosciences).

### Microinjection of mouse oocytes

In this study, mouse oocytes arrested at the GV stage were injected with different concentrations of CoA (0.15-5 mM). Since the commercial preparation of CoA used for microinjection contained 3 moles of lithium ion per mole of CoA, control oocytes were injected with equivalent amount of LiCl. In a separate study, arrested oocytes were injected with 3 mM CoA, ADP, AMP, 3’-5’-ADP and pantetheine. To examine the effect of injected compounds on spindle formation, oocytes were fixed 17 hours after GV release and stained for alpha-tubulin (green) and DNA (blue)

### Intact Mass Spectrometry

To generate CoA complexes with Aurora A, 10 µM WT Aurora A was incubated with 1 µM Coenzyme A for 15 minutes at room temperature. To evaluate the interaction, intact complexes were desalted using a C4 desalting trap (Waters MassPREP™ Micro desalting column, 2.1 × 5 mm, 20 μ m particle size, 1000 Å pore size). Aurora A was eluted with 50 % (v/v) MeCN, 0.1 % (v/v) formic acid. Intact mass analysis was performed using a Waters nano ACQUITY Ultra Performance liquid chromatography (UPLC) system coupled to a Waters SYNAPT G2, as described (Foulkes et al., 2018). Samples were eluted from a C4 trap column at a flow rate of 10 µL/min using three repeated 0-100 % acetonitrile gradients. Data was collected between 400 and 3500 *m/z* and processed using MaxEnt1 (maximum entropy software, Waters Corporation).

### Ion Mobility

IM-MS analysis was performed on a Waters Synapt G2-S*i* instrument. Aurora A was buffer exchanged into 50 mM NH_4_OAc (LC grade, Sigma) as previously described (Byrne et al., 2016). Typically, 1-3 µl of 2-5 µM sample was analyzed using borosilicate emitters (Thermo ES 387). Spraying voltage was adjusted to 1.1 −1.8 kV, sampling cone was 20 V. Pressure in the travelling wave (T-wave) ion mobility cell was 2.78 mbar (nitrogen), wave height was kept at 30 V, wave velocity at 750 m/s. In order to experimentally determine collision cross section (CCS), drift time through the T-wave mobility cell was performed using β-lactoglobulin A (Sigma L7880), avidin (Sigma A9275), transthyretin (Sigma P1742), concanavalin A (Sigma C2010) and serum albumin (Sigma P7656) according to standard protocols. The exact hard sphere scattering (EHSS) model implemented in the Mobcal software was used to calculate CCS values on the basis of X-ray structures, as described previously (Byrne et al., 2016).

### Mass spectrometry and data processing

*In vitro* CoAlated His-Aurora A or immunoprecipitated FLAG-Aurora A were digested with sequencing grade trypsin (Promega). After heat-inactivation of trypsin, CoAlated peptides were immunoprecipitated with anti-CoA antibody cross-linked to Protein G Sepharose. Immunoprecipitated peptides were treated with Nudix 7 and enriched further by an IMAC column before LC–MS/MS analysis. The resulting samples were analysed by nano-scale capillary LCMS/MS using an Ultimate U3000 UPLC System (Dionex) fitted with a 100 µm x 2 cm PepMap100 C18 nano trap column and a 75 μm × 25 cm PepMap100 C18 nano analytical column (Dionex). Peptides were eluted using an acetonitrile gradient and sprayed directly via a nano-flow electrospray ionization source into the mass spectrometer (Orbitrap Velos, Thermo Scientific). The mass spectrometer was operated in data dependent mode, using a full scan (m/z = 350-1600) in the Orbitrap analyser, with a resolution of 60,000 at m/z =400, followed by MS/MS acquisitions of the 20 most intense ions in the LTQ Velos ion trap. Maximum FTMS scan accumulation times were set at 250 ms and maximum ion trap MSn scan accumulation times were set at 200 ms. The Orbitrap measurements were internally calibrated using the lock mass of polydimethylcyclosiloxane at m/z 445.120025. Dynamic exclusion was set for 30 s with exclusion list of 500. LC-MS/MS raw data files were processed as standard samples using MaxQuant (Cox and Mann, 2008) version 1.5.2.8, which incorporates the Andromeda search engine. MaxQuant processed data was searched against Human UniProt protein database. Carbamidomethyl cysteine, Acetyl N-terminal, N-ethylmaleimide cysteine, oxidation of methionines, CoAlation of cysteine with delta mass 338, 356 and 765, were set as variable modifications. For all data sets, the default parameters in MaxQuant were used, except MS/MS tolerance, which was set at 0.6 Da and the second peptide ID was unselected.

---

**Supplementary Figure 1.**
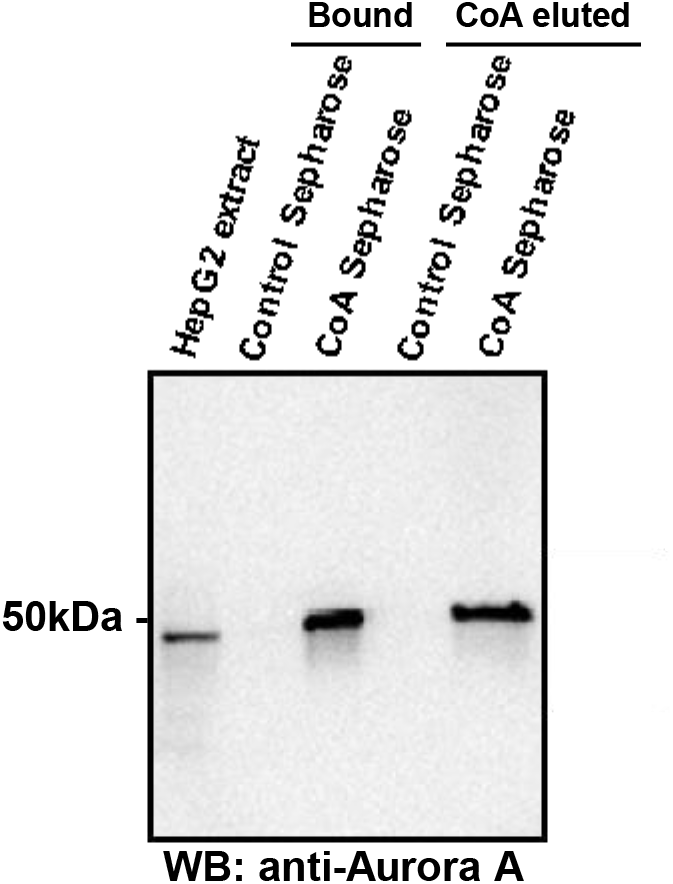
Binding of endogenous cellular Aurora A to CoA. CoA Sepharose or Sepharose control beads were employed in an immunoprecipitation assay using extracts prepared from exponentially growing HepG2 cells. Bound proteins were eluted from beads with 100 μM CoA (CoA-eluted fraction) or 2x SDS loading buffer. Eluted proteins were separated by SDS-PAGE and immunoblotted with anti-Aurora A antibody.

**Supplementary Figure 2.**
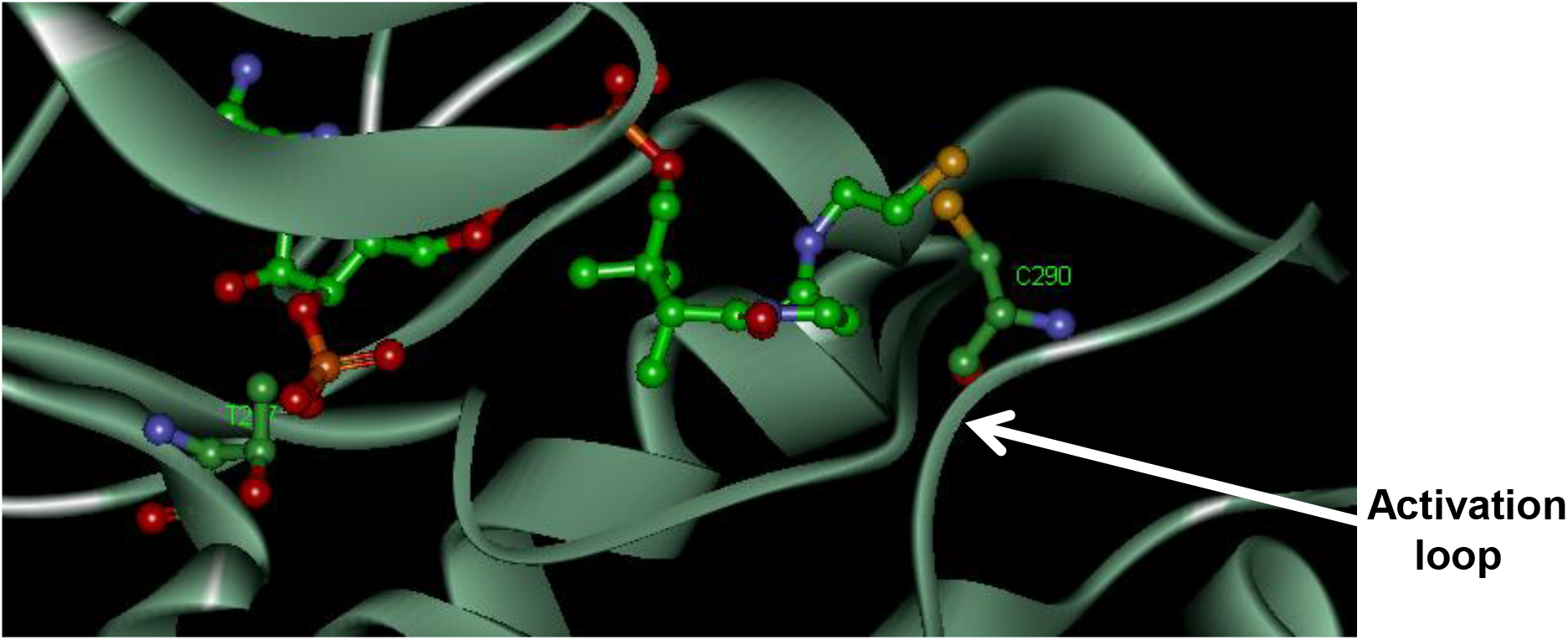
Molecular docking of CoA to Aurora A. Model of CoA bound to Aurora A (PDB 1OL7). The location of Thr 217 adjacent to the 3’phospho-adenine of CoA, and Cys 290 in the kinase activation segment, are both indicated.

**Supplementary Figure 3.**
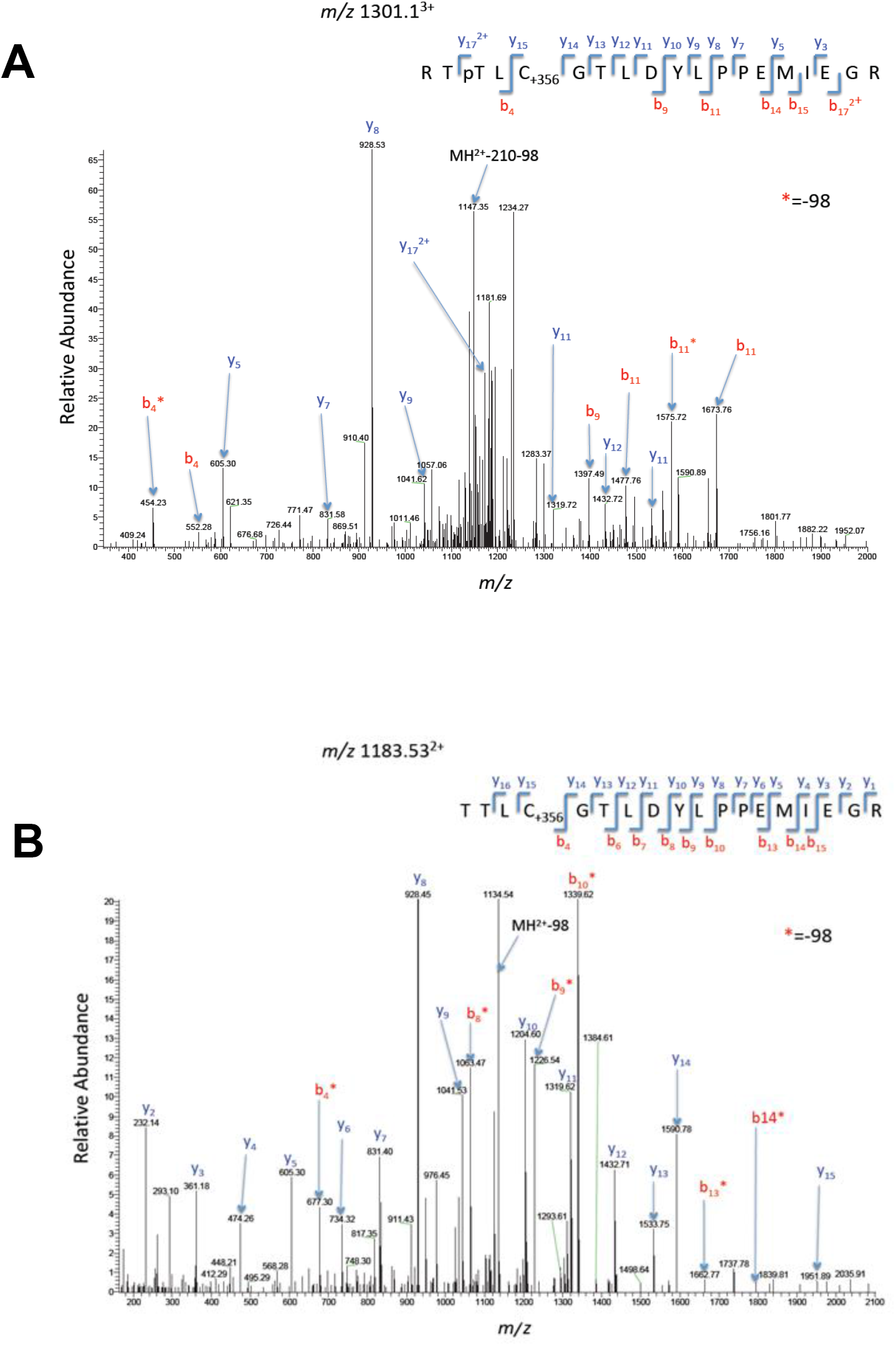
MS analysis of T288 phosphorylated and CoAlated Aurora A. LC–MS/MS spectra of Aurora A tryptic peptides derived from **(A)** *in vitro* CoAlation or **(B)** FLAG-Aurora A isolated from peroxide-treated cells, where Aurora A is CoAlated on Cys290. The ion chromatogram for peptide with m/z of 1183.53 is shown, CID confirmed Cys290 as the site of CoAlation.

**Supplementary Figure 4.**
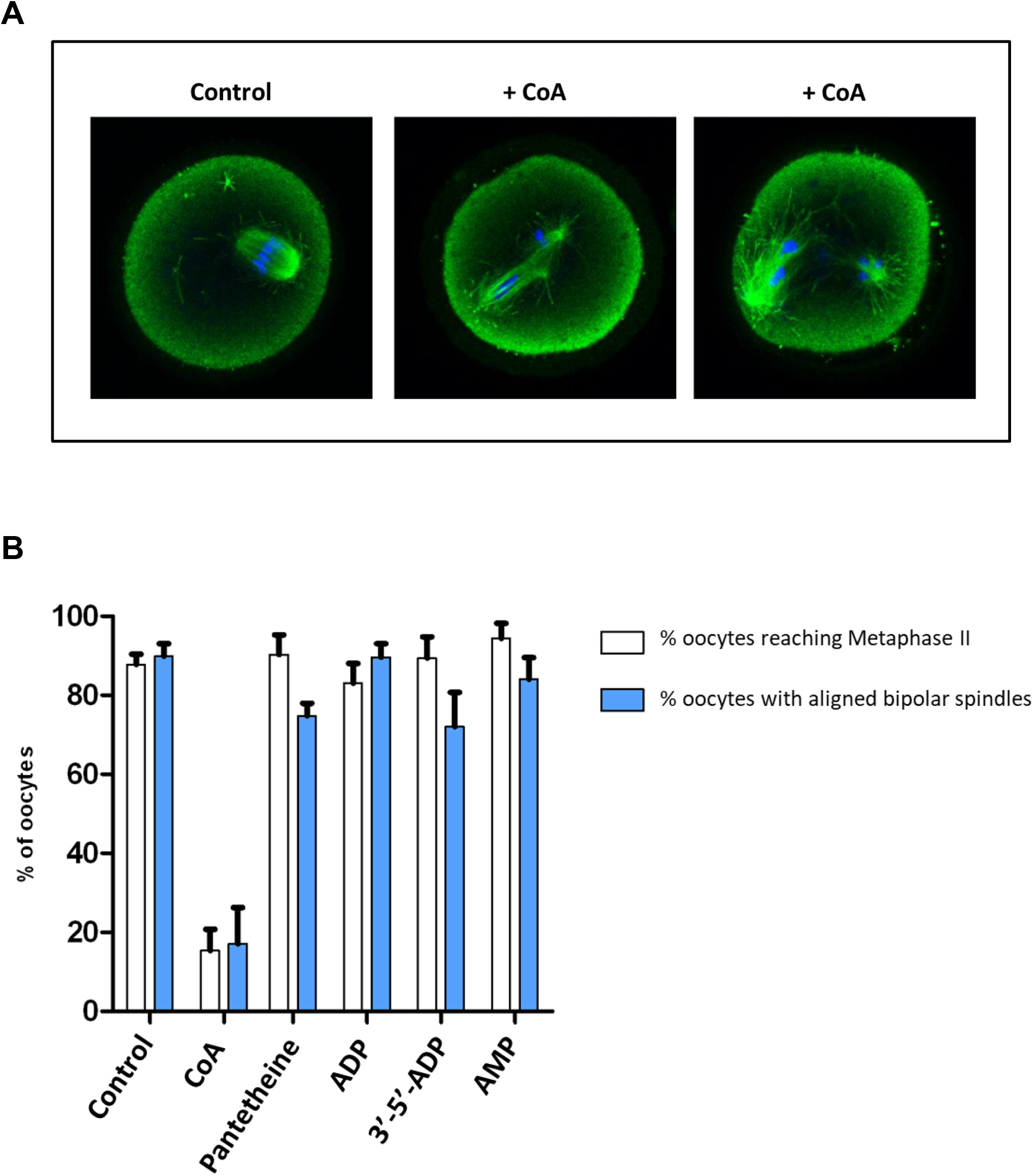
Microinjection of CoA causes abnormal spindles and chromosome misalignment in mouse oocytes. (A) Mouse oocytes arrested at the GV stage were injected with CoA or different compounds, representing integral parts of the CoA molecule. 17 hrs after GV release, cells were fixed and stained for alpha-tubulin (green) or DNA (blue), and (B) the numbers of oocytes with normal and abnormal spindles were recorded.

## References

Adams, P. D., Grosse-Kunstleve, R. W., Hung, L. W., Ioerger, T. R., Mccoy, A. J., Moriarty, N. W., Read, R. J., Sacchettini, J. C., Sauter, N. K. & Terwilliger, T. C. 2002. PHENIX: building new software for automated crystallographic structure determination. Acta crystallographica. Section D, Biological crystallography, 58, 1948-54.

Bayliss, R., Sardon, T., Ebert, J., Lindner, D., Vernos, I. & Conti, E. 2004. Determinants for Aurora-A activation and Aurora-B discrimination by TPX2. Cell cycle, 3, 404-7.

Bayliss, R., Sardon, T., Vernos, I. & Conti, E. 2003. Structural basis of Aurora-A activation by TPX2 at the mitotic spindle. Molecular cell, 12, 851-62.

Bertolin, G., Bulteau, A. L., Alves-Guerra, M. C., Burel, A., Lavault, M. T., Gavard, O., Le BRAS, S., Gagne, J. P., Poirier, G. G., Le BORGNE, R., Prigent, C. & Tramier, M. 2018. Aurora kinase A localises to mitochondria to control organelle dynamics and energy production. eLife, 7.

Bischoff, J. R., Anderson, L., Zhu, Y., Mossie, K., Ng, L., Souza, B., Schryver, B., Flanagan, P., Clairvoyant, F., Ginther, C., Chan, C. S., Novotny, M., Slamon, D. J. & Plowman, G. D. 1998. A homologue of Drosophila aurora kinase is oncogenic and amplified in human colorectal cancers. The EMBO journal, 17, 3052-65.

Breus, O., Panasyuk, G., Gout, I.T., Filonenko, V. & Nemazanyy I. 2009. CoA synthase is in complex with p85alpha PI3K and affects PI3K signaling pathway. Biochem Biophys Res Commun. 385, 581-5.

Breus, O., Panasyuk, G., Gout, I.T., Filonenko, V. & Nemazanyy, I. 2010. CoA Synthase is phosphorylated on tyrosines in mammalian cells, interacts with and is dephosphorylated by Shp2PTP. Mol Cell Biochem. 335, 195-202.

Burgess, S. G. & Bayliss, R. 2015. The structure of C290A:C393A Aurora A provides structural insights into kinase regulation. Acta crystallographica. Section F, Structural biology communications, 71, 315-9.

Burgess, S. G., Mukherjee, M., Sabir, S., Joseph, N., Gutierrez-Caballero, C., Richards, M. W., Huguenin-Dezot, N., Chin, J. W., Kennedy, E. J., Pfuhl, M., Royle, S. J., Gergely, F. & Bayliss, R. 2018. Mitotic spindle association of TACC3 requires Aurora-A-dependent stabilization of a cryptic alpha-helix. The EMBO journal, 37.

Burgess, S. G., Peset, I., Joseph, N., Cavazza, T., Vernos, I., Pfuhl, M., Gergely, F. & Bayliss, R. 2015. Aurora-A-Dependent Control of TACC3 Influences the Rate of Mitotic Spindle Assembly. PLoS genetics, 11, e1005345.

Bury, L., Coelho, P. A., Simeone, A., Ferries, S., Eyers, C. E., Eyers, P. A., Zernicka-GOETZ, M. & Glover, D. M. 2017. Plk4 and Aurora A cooperate in the initiation of acentriolar spindle assembly in mammalian oocytes. The Journal of cell biology, 216, 3571-3590.

Byrne, D. P., Li, Y., Ngamlert, P., Ramakrishnan, K., Eyers, C. E., Wells, C., Drewry, D. H., Zuercher, W. J., Berry, N. G., Fernig, D. G. & Eyers, P. A. 2018. New tools for evaluating protein tyrosine sulfation: tyrosylprotein sulfotransferases (TPSTs) are novel targets for RAF protein kinase inhibitors. The Biochemical journal, 475, 2435-2455.

Byrne, D. P., Vonderach, M., Ferries, S., Brownridge, P. J., Eyers, C. E. & Eyers, P. A. 2016. cAMP-dependent protein kinase (PKA) complexes probed by complementary differential scanning fluorimetry and ion mobility-mass spectrometry. The Biochemical journal, 473, 3159-75.

Carmena, M. & Earnshaw, W. C. 2003. The cellular geography of aurora kinases. Nature reviews. Molecular cell biology, 4, 842-54.

Carmena, M., Ruchaud, S. & Earnshaw, W. C. 2009. Making the Auroras glow: regulation of Aurora A and B kinase function by interacting proteins. Current opinion in cell biology, 21, 796-805.

Chen, V. B., Arendall, W. B., 3RD, Headd, J. J., Keedy, D. A., Immormino, R. M., Kapral, G. J., Murray, L. W., Richardson, J. S. & Richardson, D. C. 2010. MolProbity: all-atom structure validation for macromolecular crystallography. Acta crystallographica. Section D, Biological crystallography, 66, 12-21.

Chiu, J. & Dawes, I. W. 2012. Redox control of cell proliferation. Trends in cell biology, 22, 592-601.

Corcoran, A. & Cotter, T. G. 2013. Redox regulation of protein kinases. The FEBS journal, 280, 1944-65.

Courtheoux, T., Diallo, A., Damodaran, A. P., Reboutier, D., Watrin, E. & Prigent, C. 2018. Aurora A kinase activity is required to maintain an active spindle assembly checkpoint during prometaphase. Journal of cell science, 131.

Cox, J. & Mann, M. 2008. MaxQuant enables high peptide identification rates, individualized p.p.b.-range mass accuracies and proteome-wide protein quantification. Nature biotechnology, 26, 1367-72.

D’ASSORO, A. B., Haddad, T. & Galanis, E. 2015. Aurora-A Kinase as a Promising Therapeutic Target in Cancer. Frontiers in oncology, 5, 295.

Damodaran, A. P., Vaufrey, L., Gavard, O. & Prigent, C. 2017. Aurora A Kinase Is a Priority Pharmaceutical Target for the Treatment of Cancers. Trends in pharmacological sciences, 38, 687-700.

Dansie, L.E., Reeves, S., Miller, K., Zano, S.P., Frank, M., Pate, C., Wang, J. & Jackowski, S. 2014. Physiological roles of the pantothenate kinases. Biochem Soc Trans. 42, 1033-6.

Dodson, C. A. & Bayliss, R. 2012. Activation of Aurora-A kinase by protein partner binding and phosphorylation are independent and synergistic. The Journal of biological chemistry, 287, 1150-7.

Dodson, C. A., Kosmopoulou, M., Richards, M. W., Atrash, B., Bavetsias, V., Blagg, J. & Bayliss, R. 2010. Crystal structure of an Aurora-A mutant that mimics Aurora-B bound to MLN8054: insights into selectivity and drug design. The Biochemical journal, 427, 19-28.

Dodson, C. A., Yeoh, S., Haq, T. & Bayliss, R. 2013. A kinetic test characterizes kinase intramolecular and intermolecular autophosphorylation mechanisms. Science signaling, 6, ra54.

Dusi, S., Valletta, L., Haack, T.B., Tsuchiya, Y., Venco, P., Pasqualato, S., Goffrini, P., Tigano, M., Demchenko, N., Wieland, T., Schwarzmayr, T., Strom, T.M., Invernizzi, F., Garavaglia, B., Gregory, A., Sanford, L., Hamada, J., Bettencourt, C., Houlden, H., Chiapparini, L., Zorzi, G., Kurian, M.A., Nardocci, N., Prokisch, H., Hayflick, S., Gout, I. & Tiranti V. 2014. Exome sequence reveals mutations in CoA synthase as a cause of neurodegeneration with brain iron accumulation. Am J Hum Genet. 94, 11-22.

Eckerdt, F., Eyers, P. A., Lewellyn, A. L., Prigent, C. & Maller, J. L. 2008. Spindle pole regulation by a discrete Eg5-interacting domain in TPX2. Current biology : CB, 18, 519-25.

Emsley, P. & Cowtan, K. 2004. Coot: model-building tools for molecular graphics. Acta crystallographica. Section D, Biological crystallography, 60, 2126-32.

Eyers, P. A., Churchill, M. E. & Maller, J. L. 2005. The Aurora A and Aurora B protein kinases: a single amino acid difference controls intrinsic activity and activation by TPX2. Cell cycle, 4, 784-9.

Eyers, P. A., Erikson, E., Chen, L. G. & Maller, J. L. 2003. A novel mechanism for activation of the protein kinase Aurora A. Current biology : CB, 13, 691-7.

Foulkes, D. M., Byrne, D. P., Yeung, W., Shrestha, S., Bailey, F. P., Ferries, S., Eyers, C. E., Keeshan, K., Wells, C., Drewry, D. H., Zuercher, W. J., Kannan, N. & Eyers, P. A. 2018. Covalent inhibitors of EGFR family protein kinases induce degradation of human Tribbles 2 (TRIB2) pseudokinase in cancer cells. Science signaling, 11.

Fu, J., Bian, M., Liu, J., Jiang, Q. & Zhang, C. 2009. A single amino acid change converts Aurora-A into Aurora-B-like kinase in terms of partner specificity and cellular function. Proceedings of the National Academy of Sciences of the United States of America, 106, 6939-44.

Girdler, F., Gascoigne, K. E., Eyers, P. A., Hartmuth, S., Crafter, C., Foote, K. M., Keen, N. J. & Taylor, S. S. 2006. Validating Aurora B as an anti-cancer drug target. Journal of cell science, 119, 3664-75.

Goos, J. A., Coupe, V. M., Diosdado, B., Delis-Van Diemen, P. M., Karga, C., Belien, J. A., Carvalho, B., Van Den Tol, M. P., Verheul, H. M., Geldof, A. A., Meijer, G. A., Hoekstra, O. S. & Fijneman, R. J. 2013. Aurora kinase A (AURKA) expression in colorectal cancer liver metastasis is associated with poor prognosis. British journal of cancer, 109, 2445-52.

Gout, I. 2018. Coenzyme A, protein CoAlation and redox regulation in mammalian cells. Biochemical Society transactions, 46, 721-728.

Gruss, O. J., Wittmann, M., Yokoyama, H., Pepperkok, R., Kufer, T., Sillje, H., Karsenti, E., Mattaj, I. W. & Vernos, I. 2002. Chromosome-induced microtubule assembly mediated by TPX2 is required for spindle formation in HeLa cells. Nature cell biology, 4, 871-9.

Gudkova, D., Panasyuk, G., Nemazanyy, I., Zhyvoloup, A., Monteil, P., Filonenko, V & Gout, I. 2012. EDC4 interacts with and regulates the dephospho-CoA kinase activity of CoA synthase. FEBS Lett. 586, 3590-5.

Haydon, C. E., Eyers, P. A., Aveline-Wolf, L. D., Resing, K. A., Maller, J. L. & Ahn, N. G. 2003. Identification of novel phosphorylation sites on Xenopus laevis Aurora A and analysis of phosphopeptide enrichment by immobilized metal-affinity chromatography. Molecular & cellular proteomics : MCP, 2, 1055-67.

Hegarat, N., Smith, E., Nayak, G., Takeda, S., Eyers, P. A. & Hochegger, H. 2011. Aurora A and Aurora B jointly coordinate chromosome segregation and anaphase microtubule dynamics. The Journal of cell biology, 195, 1103-13.

Humphries, K. M., Deal, M. S. & Taylor, S. S. 2005. Enhanced dephosphorylation of cAMP-dependent protein kinase by oxidation and thiol modification. The Journal of biological chemistry, 280, 2750-8.

Humphries, K. M., Juliano, C. & Taylor, S. S. 2002. Regulation of cAMP-dependent protein kinase activity by glutathionylation. The Journal of biological chemistry, 277, 43505-11.

Humphries, K. M., Pennypacker, J. K. & Taylor, S. S. 2007. Redox regulation of cAMP-dependent protein kinase signaling: kinase versus phosphatase inactivation. The Journal of biological chemistry, 282, 22072-9.

Joukov, V. & De Nicolo, A. 2018. Aurora-PLK1 cascades as key signaling modules in the regulation of mitosis. Science signaling, 11.

Keen, N. & Taylor, S. 2004. Aurora-kinase inhibitors as anticancer agents. Nature reviews. Cancer, 4, 927-36.

Kufer, T. A., Sillje, H. H., Korner, R., Gruss, O. J., Meraldi, P. & Nigg, E. A. 2002. Human TPX2 is required for targeting Aurora-A kinase to the spindle. The Journal of cell biology, 158, 617-23.

Leonardi, R., Zhang, Y.M., Rock, C.O. & Jackowski, S. 2005. Coenzyme A: back in action. Prog Lipid Res. 44, 125-53

Levinson, N. M. 2018. The multifaceted allosteric regulation of Aurora kinase A. The Biochemical journal, 475, 2025-2042.

Macurek, L., Lindqvist, A., Lim, D., Lampson, M. A., Klompmaker, R., Freire, R., Clouin, C., Taylor, S. S., Yaffe, M. B. & Medema, R. H. 2008. Polo-like kinase-1 is activated by aurora A to promote checkpoint recovery. Nature, 455, 119-23.

Malanchuk, O.M., Panasyuk, G.G., Serbyn, N.M., Gout, I.T. & Filonenko, V.V. 2015. Generation and characterization of monoclonal antibodies specific to coenzyme A. Biopolym. Cell. 31, 187–19

Manfredi, M. G., Ecsedy, J. A., Meetze, K. A., Balani, S. K., Burenkova, O., Chen, W., Galvin, K. M., Hoar, K. M., Huck, J. J., Leroy, P. J., Ray, E. T., Sells, T. B., Stringer, B., Stroud, S. G., Vos, T. J., Weatherhead, G. S., Wysong, D. R., Zhang, M., Bolen, J. B. & Claiborne, C. F. 2007. Antitumor activity of MLN8054, an orally active small-molecule inhibitor of Aurora A kinase. Proceedings of the National Academy of Sciences of the United States of America, 104, 4106-11.

Martinez, D.L., Tsuchiya, Y. & Gout, I. 2014. Coenzyme A biosynthetic machinery in mammalian cells. Biochem Soc Trans. 42, 1112-7.

Mccoy, A. J., Grosse-Kunstleve, R. W., Adams, P. D., Winn, M. D., Storoni, L. C. & Read, R. J. 2007. Phaser crystallographic software. Journal of applied crystallography, 40, 658-674.

Mcintyre, P. J., Collins, P. M., Vrzal, L., Birchall, K., Arnold, L. H., Mpamhanga, C., Coombs, P. J., Burgess, S. G., Richards, M. W., Winter, A., Veverka, V., Delft, F. V., Merritt, A. & Bayliss, R. 2017. Characterization of Three Druggable Hot-Spots in the Aurora-A/TPX2 Interaction Using Biochemical, Biophysical, and Fragment-Based Approaches. ACS chemical biology, 12, 2906-2914.

Murphy, J. M., Zhang, Q., Young, S. N., Reese, M. L., Bailey, F. P., Eyers, P. A., Ungureanu, D., Hammaren, H., Silvennoinen, O., Varghese, L. N., Chen, K., Tripaydonis, A., Jura, N., Fukuda, K., Qin, J., Nimchuk, Z., Mudgett, M. B., Elowe, S., Gee, C. L., Liu, L., Daly, R. J., Manning, G., Babon, J. J. & Lucet, I. S. 2014. A robust methodology to subclassify pseudokinases based on their nucleotide-binding properties. The Biochemical journal, 457, 323-34.

Ocasio, C. A., Warkentin, A. A., Mcintyre, P. J., Barkovich, K. J., Vesely, C., Spencer, J., Shokat, K. M. & Bayliss, R. 2018. Type II kinase inhibitors targeting Cys-gatekeeper kinases display orthogonality with wild type and Ala/Gly-gatekeeper kinases. ACS chemical biology.

Ohashi, S., Sakashita, G., Ban, R., Nagasawa, M., Matsuzaki, H., Murata, Y., Taniguchi, H., Shima, H., Furukawa, K. & Urano, T. 2006. Phospho-regulation of human protein kinase Aurora-A: analysis using anti-phospho-Thr288 monoclonal antibodies. Oncogene, 25, 7691-702.

Ostrem, J. M., Peters, U., Sos, M. L., Wells, J. A. & Shokat, K. M. 2013. K-Ras(G12C) inhibitors allosterically control GTP affinity and effector interactions. Nature, 503, 548-51.

Pitsawong, W., Buosi, V., Otten, R., Agafonov, R. V., Zorba, A., Kern, N., Kutter, S., Kern, G., Padua, R. A., Meniche, X. & Kern, D. 2018. Dynamics of human protein kinase Aurora A linked to drug selectivity. eLife, 7.

Reibel, D.K., Wyse, B.W., Berkich, D.A. & Neely, J.R. 1981. Regulation of coenzyme A synthesis in heart muscle: effects of diabetes and fasting. Am J Physiol. 240, H606-11.

Richards, M. W., Burgess, S. G., Poon, E., Carstensen, A., Eilers, M., Chesler, L. & Bayliss, R. 2016. Structural basis of N-Myc binding by Aurora-A and its destabilization by kinase inhibitors. Proceedings of the National Academy of Sciences of the United States of America, 113, 13726-13731.

Rock, C.O., Calder, R.B., Karim, M.A. & Jackowski, S. 2000. Pantothenate kinase regulation of the intracellular concentration of coenzyme A. J Biol Chem. 275, 1377-83.

Sardon, T., Pache, R. A., Stein, A., Molina, H., Vernos, I. & Aloy, P. 2010. Uncovering new substrates for Aurora A kinase. EMBO reports, 11, 977-84.

Saskova, A., Solc, P., Baran, V., Kubelka, M., Schultz, R. M. & Motlik, J. 2008. Aurora kinase A controls meiosis I progression in mouse oocytes. Cell cycle, 7, 2368-76.

Savitsky, P. A. & Finkel, T. 2002. Redox regulation of Cdc25C. The Journal of biological chemistry, 277, 20535-40.

Scutt, P. J., Chu, M. L., Sloane, D. A., Cherry, M., Bignell, C. R., Williams, D. H. & Eyers, P. A. 2009. Discovery and exploitation of inhibitor-resistant aurora and polo kinase mutants for the analysis of mitotic networks. The Journal of biological chemistry, 284, 15880-93.

Sibon, O.C. & Strauss, E. 2016. Coenzyme A: to make it or uptake it? Nat Rev Mol Cell Biol. 17, 605-606.

Sloane, D. A., Trikic, M. Z., Chu, M. L., Lamers, M. B., Mason, C. S., Mueller, I., Savory, W. J., Williams, D. H. & Eyers, P. A. 2010. Drug-resistant aurora A mutants for cellular target validation of the small molecule kinase inhibitors MLN8054 and MLN8237. ACS chemical biology, 5, 563-76.

Tayyar, Y., Jubair, L., Fallaha, S. & Mcmillan, N. A. J. 2017. Critical risk-benefit assessment of the novel anti-cancer aurora a kinase inhibitor alisertib (MLN8237): A comprehensive review of the clinical data. Critical reviews in oncology/hematology, 119, 59-65.

Tonks, N. K. 2005. Redox redux: revisiting PTPs and the control of cell signaling. Cell, 121, 667-70.

Tsuchiya, Y., Peak-CHEW, S. Y., Newell, C., Miller-AIDOO, S., Mangal, S., Zhyvoloup, A., Bakovic, J., Malanchuk, O., Pereira, G. C., Kotiadis, V., Szabadkai, G., Duchen, M. R., Campbell, M., Cuenca, S. R., Vidal-PUIG, A., James, A. M., Murphy, M. P., Filonenko, V., Skehel, M. & Gout, I. 2017. Protein CoAlation: a redox-regulated protein modification by coenzyme A in mammalian cells. The Biochemical journal, 474, 2489-2508.

Tsuchiya, Y., Pham, U., Hu, W., Ohnuma, S. & Gout, I. 2014. Changes in acetyl CoA levels during the early embryonic development of Xenopus laevis. PloS one, 9, e97693.

Tsuchiya, Y., Zhyvoloup, A., Bakovic, J., Thomas, N., Yu, B. Y. K., Das, S., Orengo, C., Newell, C., Ward, J., Saladino, G., Comitani, F., Gervasio, F. L., Malanchuk, O. M., Khoruzhenko, A. I., Filonenko, V., Peak-CHEW, S. Y., Skehel, M. & Gout, I. 2018. Protein CoAlation and antioxidant function of coenzyme A in prokaryotic cells. The Biochemical journal, 475, 1909-1937.

Tyler, R. K., Shpiro, N., Marquez, R. & Eyers, P. A. 2007. VX-680 inhibits Aurora A and Aurora B kinase activity in human cells. Cell cycle, 6, 2846-54.

Vo, T. T. L., Park, J. H., Seo, J. H., Lee, E. J., Choi, H., Bae, S. J., Le, H., An, S., Lee, H. S., Wee, H. J. & Kim, K. W. 2017. ARD1-mediated aurora kinase A acetylation promotes cell proliferation and migration. Oncotarget, 8, 57216-57230.

Walter, A. O., Seghezzi, W., Korver, W., Sheung, J. & Lees, E. 2000. The mitotic serine/threonine kinase Aurora2/AIK is regulated by phosphorylation and degradation. Oncogene, 19, 4906-16.

Wang, G. F., Dong, Q., Bai, Y., Yuan, J., Xu, Q., Cao, C. & Liu, X. 2017. Oxidative stress induces mitotic arrest by inhibiting Aurora A-involved mitotic spindle formation. Free radical biology & medicine, 103, 177-187.

Winter, G., Lobley, C. M. & Prince, S. M. 2013. Decision making in xia2. Acta crystallographica. Section D, Biological crystallography, 69, 1260-73.

Wittmann, T., Wilm, M., Karsenti, E. & Vernos, I. 2000. TPX2, A novel xenopus MAP involved in spindle pole organization. The Journal of cell biology, 149, 1405-18.

Yasui, Y., Urano, T., Kawajiri, A., Nagata, K., Tatsuka, M., Saya, H., Furukawa, K., Takahashi, T., Izawa, I. & Inagaki, M. 2004. Autophosphorylation of a newly identified site of Aurora-B is indispensable for cytokinesis. The Journal of biological chemistry, 279, 12997-3003.

Yu, X., Minter-Dykhouse, K., Malureanu, L., Zhao, W. M., Zhang, D., Merkle, C. J., Ward, I. M., Saya, H., Fang, G., Van Deursen, J. & Chen, J. 2005. Chfr is required for tumor suppression and Aurora A regulation. Nature genetics, 37, 401-6.

Zeng, K., Bastos, R. N., Barr, F. A. & Gruneberg, U. 2010. Protein phosphatase 6 regulates mitotic spindle formation by controlling the T-loop phosphorylation state of Aurora A bound to its activator TPX2. The Journal of cell biology, 191, 1315-32.

Zhang, T., Kwiatkowski, N., Olson, C. M., Dixon-Clarke, S. E., Abraham, B. J., Greifenberg, A. K., Ficarro, S. B., Elkins, J. M., Liang, Y., Hannett, N. M., Manz, T., Hao, M., Bartkowiak, B., Greenleaf, A. L., Marto, J. A., Geyer, M., Bullock, A. N., Young, R. A. & Gray, N. S. 2016. Covalent targeting of remote cysteine residues to develop CDK12 and CDK13 inhibitors. Nature chemical biology, 12, 876-84.

Zhang, Y.M., Rock, C.O. & Jackowski, S. 2005. Feedback regulation of murine pantothenate kinase 3 by coenzyme A and coenzyme A thioesters. J Biol Chem., 280, 32594-601.

Zhao, Z. & Bourne, P. E. 2018. Progress with covalent small-molecule kinase inhibitors. Drug discovery today, 23, 727-735.

Zhao, Z., Liu, Q., Bliven, S., Xie, L. & Bourne, P. E. 2017. Determining Cysteines Available for Covalent Inhibition Across the Human Kinome. Journal of medicinal chemistry, 60, 2879-2889.

Zhou, B., Westaway, S.K., Levinson, B., Johnson, M.A., Gitschier, J. & Hayflick, S.J. 2001. A novel pantothenate kinase gene (PANK2) is defective in Hallervorden-Spatz syndrome. Nat Genet. 2001. 28, 345-9.

Zhou, H., Kuang, J., Zhong, L., Kuo, W. L., Gray, J. W., Sahin, A., Brinkley, B. R. & Sen, S. 1998. Tumour amplified kinase STK15/BTAK induces centrosome amplification, aneuploidy and transformation. Nature genetics, 20, 189-93.

Zhyvoloup, A., Nemazanyy, I., Panasyuk, G., Valovka, T., Fenton, T., Rebholz, H., Wang, M.L., Foxon, R., Lyzogubov, V., Usenko, V., Kyyamova, R., Gorbenko, O., Matsuka, G., Filonenko, V. & Gout, I.T. 2003. Subcellular localization and regulation of coenzyme A synthase. J Biol Chem. 278, 50316-21.

